# The critical role of the ZBP1-NINJ1 axis and IRF1/IRF9 in ethanol-induced cell death, PANoptosis, and alcohol-associated liver disease

**DOI:** 10.1101/2025.03.12.642836

**Authors:** Qiang Qin, Wen Chen, Clay D. King, Sivakumar Prasanth Kumar, Peter Vogel, Rebecca E. Tweedell, Thirumala-Devi Kanneganti

## Abstract

Innate immunity provides the critical first line of defense against infection and sterile triggers. Cell death is a key component of the innate immune response to clear pathogens, but excessive or aberrant cell death can induce inflammation, cytokine storm, and pathology, making it a central molecular mechanism in inflammatory diseases. Alcohol-associated liver disease (ALD) is one such inflammatory disease, but the specific innate immune mechanisms driving pathology in this context remain unclear. Here, by leveraging RNAseq and tissue expression in clinical samples, we identified increased expression of the innate immune sensor Z-DNA binding protein (ZBP1) in patients with ALD. We discovered that ZBP1 expression correlated with ALD progression in patients, and that ethanol induced ZBP1-dependent lytic cell death, PANoptosis, in immune (macrophages, monocytes, Kupffer cells) and non-immune cells (hepatocytes). Mechanistically, the interferon regulatory factors (IRFs) IRF9 and IRF1 upregulated ZBP1 expression, allowing ZBP1 to sense Z-NAs through its Zα2 domain and drive PANoptosis signaling, cell membrane rupture through NINJ1, and DAMP release. Furthermore, the expressions of ZBP1 and NINJ1 were upregulated in both liver and serum samples from patients with ALD. In mouse models of chronic and acute ALD, ZBP1-deficient mice were significantly protected from disease pathology and liver damage. Overall, our findings establish the critical role of the ZBP1-NINJ1 axis regulated by IRFs in driving inflammatory cell death, PANoptosis, in liver cells, suggesting that targeting these molecules will have therapeutic potential in ALD and other inflammatory conditions.

## Introduction

The innate immune system serves as the first line of defense against infections and sterile insults. Upon detection of pathogens, pathogen-associated molecular patterns (PAMPs), and damage-associated molecular patterns (DAMPs), pattern recognition receptors (PRRs) activate immune signaling cascades to drive inflammation and cell death.^1^ While cell death is essential for host defense, aberrant cell death can induce pathological inflammation, leading to inflammatory diseases. Alcohol-associated liver disease (ALD) represents one such family of inflammatory diseases, and it is the leading cause of liver-related morbidity and mortality worldwide.^2^ ALD encompasses a set of pathological conditions that are caused by chronic alcohol consumption. Key roles for innate immune activation and inflammation have been identified in driving ALD pathogenesis.^3–8^ At the cellular level, alcohol exposure elicits proinflammatory responses from immune cells,^9^ subsequently inducing liver inflammation, as well as cellular and tissue damage, which can ultimately progress to ALD.^10^ Although cell death has been associated with ALD,^11^ the underlying molecular mechanisms of innate immune activation driving this pathology remain poorly understood. This knowledge gap limits the development of pharmacological interventions, and currently, there are no FDA-approved therapies for ALD. Therefore, understanding the fundamental innate immune responses that drive inflammation is critical to identify new therapeutic targets.

In this study, we sought to understand the roles of innate immune sensors and cell death pathways in inflammation and ALD pathogenesis. Our study identified ZBP1 as a key innate immune sensor in patients with ALD, as its expression correlated with disease progression. We also discovered that ethanol drove ZBP1-dependent inflammatory cell death in immune and non-immune cells through the ZBP1-NINJ1 axis, with the expression of ZBP1 and NINJ1 being upregulated in both liver and serum samples from patients with ALD. Mechanistically, interferon regulatory factors IRF9 and IRF1 regulated ZBP1 expression. ZBP1 then utilized its Zα2 domain to sense Z-NAs, inducing PANoptosis and NINJ1-mediated cell membrane rupture. Furthermore, ZBP1-deficient mice were protected from disease in both chronic and acute ALD models. Together, our findings identify a distinct inflammatory pathway driven by ethanol as an innate immune stimulus, leading to cell death through the ZBP1-NINJ1 axis downstream of IRF9/IRF1. These findings suggest that ZBP1 and NINJ1 may serve as biomarkers for ALD progression, and that targeting these molecules may mitigate inflammation and serve as prospective therapeutics for ALD and inflammatory conditions more broadly.

## Results

### ZBP1 expression is upregulated in patients with different stages of ALD

ALD is a family of inflammatory conditions,^11^ but the specific innate immune sensors and mechanisms involved in this process remain poorly characterized, making it difficult to develop targeted therapeutic strategies. Therefore, we sought to understand the molecular mechanisms of disease in ALD. We first examined patient samples from varying stages of ALD (ALD progresses through three histological stages: i) steatosis or fatty liver, characterized by fat accumulation in the liver parenchyma; ii) alcoholic hepatitis (AH), marked with steatosis, hepatocellular necrosis, and acute inflammation in liver tissues; and iii) alcoholic cirrhosis (AC), a stage of irreversible damage with fibrotic septage surrounding regenerative nodules in the liver).^12,13^ Severe AH and AC are associated with a high risk of hepatocellular carcinoma and mortality. To identify the role of innate immune sensors in ALD, we analyzed RNA sequencing data from liver tissues of healthy individuals and patients with ALD (GSE143318).^14,15^ Expression of several innate immune nucleic acid sensors was significantly upregulated in patients with ALD compared to the healthy controls, including Z-DNA binding protein 1 (*ZBP1*), interferon (IFN)-γ– inducible protein 16 (*IFI16*), retinoic acid inducible gene I receptor (*RIGI*), absent in melanoma 2 (*AIM2*), and cyclic GMP-AMP synthase (*CGAS*) and its downstream interactor stimulator of IFN genes (*STING*) (**Figure 1A**). Among these, *ZBP1* was the most significantly upregulated gene in patients with ALD (**Figure 1A and S1A**). Additionally, ZBP1 protein levels were increased in liver tissue from patients with ALD, with a moderate increase in patients with alcohol-associated steatosis (AS) and a more pronounced increase in those with AC when compared to the healthy controls (**Figure 1B and 1C**). Together, RNAseq and protein expression analyses of human liver tissues suggest that the innate immune sensor ZBP1 is associated with ALD and its progression. To further understand the role of ZBP1 in alcohol-associated disease, we conducted a murine chronic-plus-binge ethanol diet model (**Figure 1D**). We found that ZBP1 protein and RNA expression levels were significantly upregulated in mice on an ethanol diet compared to the control group (**Figure 1E and 1F**). These results further suggested that ZBP1 may play a role in the development of ALD.

**Figure 1.**
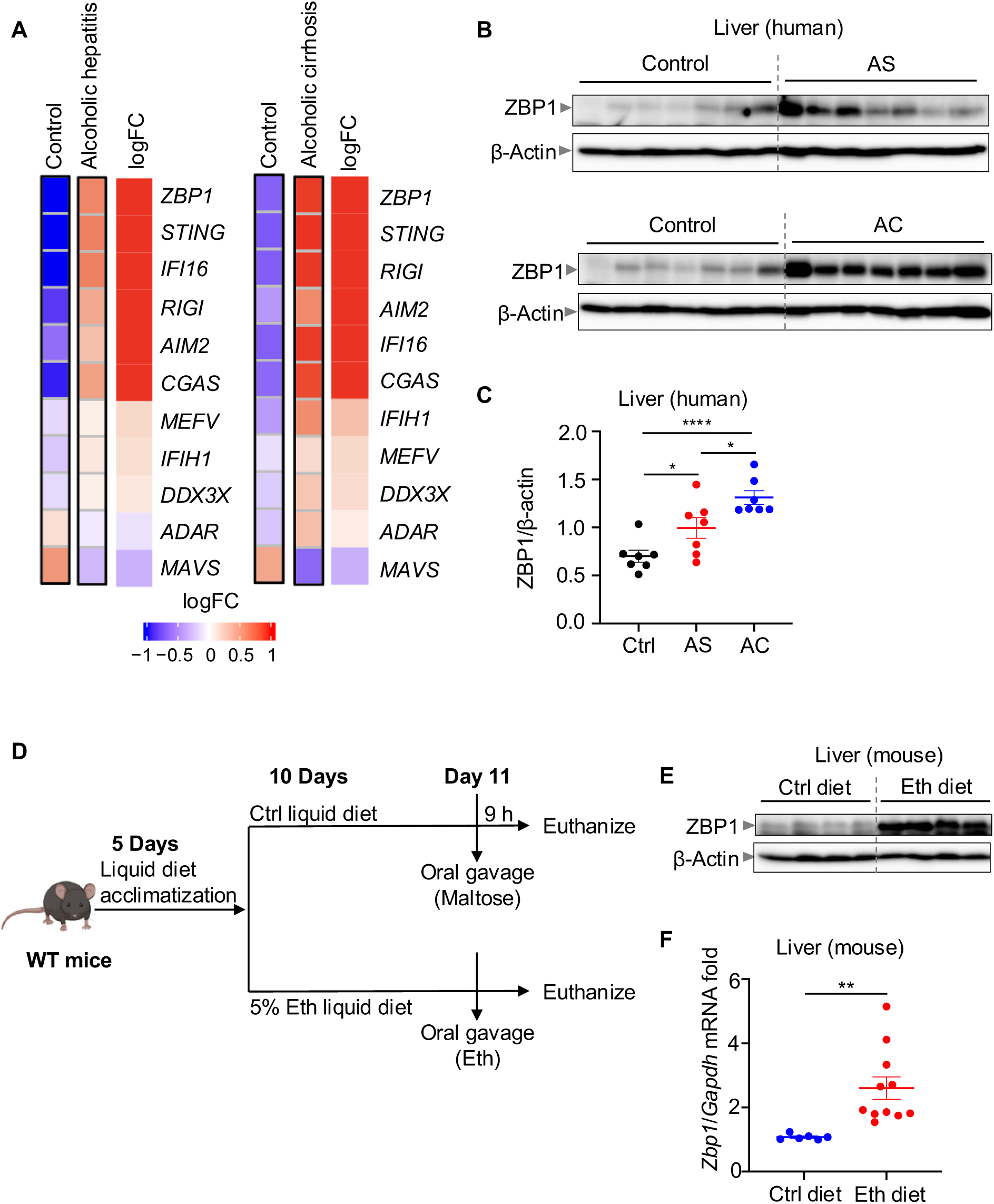
ZBP1 expression is upregulated in patients with alcohol-associated liver disease (ALD) (**A**) Heatmap depicting the expression profile of nucleic acid sensors in the liver tissues from individuals without alcohol-associated liver disease (control), patients with alcoholic hepatitis (AH), or patients with alcoholic cirrhosis (AC). The heatmap represents differentially regulated genes, where blue indicates downregulated genes and red indicates upregulated genes. (**B**) Immunoblot analysis of ZBP1 in the liver tissues from individuals without alcohol-associated liver disease (control), patients with AS, or patients with AC. β-Actin was used as the loading control. (**C**) Densitometry analysis of ZBP1 protein levels relative to β-Actin expression from liver tissues from controls and patients with AS or AC. (**D**) Schematic illustrating the chronic-plus-binge ethanol diet-induced liver disease model. (**E**) Immunoblot analysis of ZBP1 expression in the liver tissues from control (Ctrl diet) and ethanol-fed (Eth diet) groups. β-Actin was used as the internal control. (**F**) RT-PCR analysis of *Zbp1* expression relative to *Gapdh* in the liver tissues from Ctrl diet and Eth diet groups. The data are representative of two independent experiments in (**E and F**). The data are represented as the mean ± SEM in (**C and F**). The analysis was performed using the unpaired t test (one-way ANOVA) in (**C and F**). \**P* < 0.05, \*\**P* < 0.01, and \*\*\*\**P* < 0.0001.

### Ethanol induces ZBP1-dependent inflammatory cell death, PANoptosis

ZBP1 is an IFN-inducible innate immune sensor that drives inflammatory cell death, PANoptosis, in response to influenza A virus (IAV) infection,^16^ as well as in response to sterile triggers, such as nuclear export inhibitors (NEIs) plus IFN.^17^ PANoptosis is an innate immune, lytic cell death pathway initiated by innate immune sensors and driven by caspases and receptor-interacting protein kinases (RIPKs) through PANoptosomes.^18–24^ While ethanol is known to induce cell death in hepatocytes and macrophages,^8,25^ the specific mechanisms involved in ethanol-induced cell death remain unknown. Additionally, the contribution of cell death to inflammation and tissue damage in disease pathology remains unclear. Therefore, we investigated the mechanisms of ethanol-induced cell death in diverse cell types, including human hepatocyte (HepG2) and monocyte (THP-1) cells and murine primary hepatocytes, Kupffer cells, and bone marrow-derived macrophages (BMDMs). We observed that ethanol treatment resulted in the dose-dependent activation of lytic cell death in both human (**Figure S2A–S2D**) and murine cells (**Figure S3A–S3F**). To understand the molecular mechanisms of ethanol-induced cell death, we assessed the activation of different cell death proteins in BMDMs following ethanol treatment. We observed that cell death was accompanied by the activation of molecules associated with PANoptosis, including the activation and cleavage of gasdermin E (GSDME), caspases-8, -3, and -7, as well as phosphorylation of MLKL (**Figure 2A**). The P20 form of gasdermin D (GSDMD), which is an inactivated form that is induced through cleavage by apoptotic caspase-3 rather than caspase-1,^26,27^ was also induced (**Figure 2A**). Overall, these data suggest that PANoptosis may be activated in response to ethanol.

**Figure 2.**
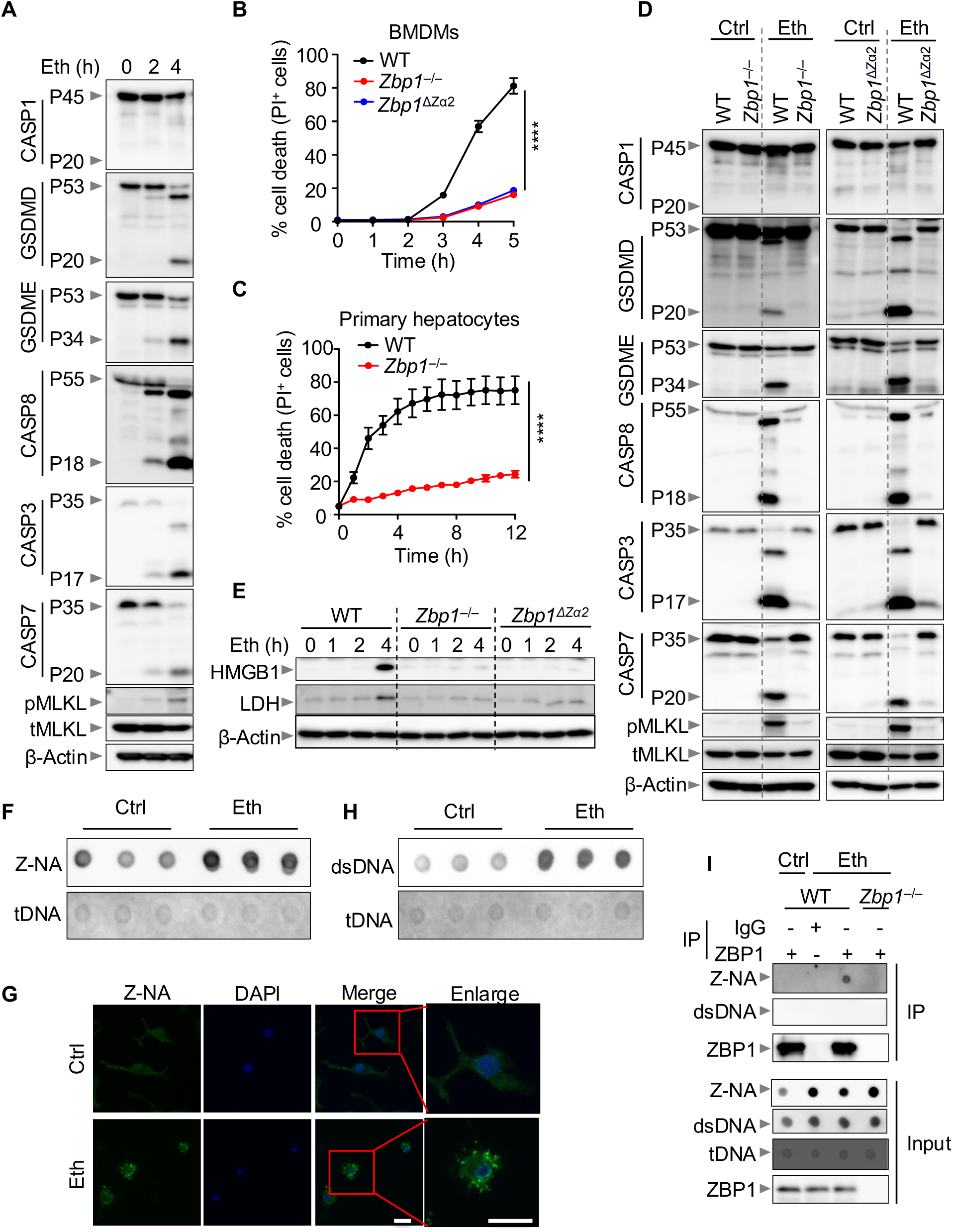
Ethanol induces ZBP1-dependent inflammatory cell death, PANoptosis. (**A**) Immunoblot analysis of pro- (P45) and activated (P20) caspase-1 (CASP1); pro- (P53) and inactivated (P20) gasdermin D (GSDMD); pro- (P53) and activated (P34) gasdermin E (GSDME); pro- (P55) and cleaved (P18) caspase-8 (CASP8); pro- (P35) and cleaved (P17) caspase-3 (CASP3); pro- (P35) and cleaved (P20) caspase-7 (CASP7); phosphorylated mixed lineage kinase domain-like pseudokinase (pMLKL), and total MLKL (tMLKL) in wild type (WT) bone marrow-derived macrophages (BMDMs) stimulated with ethanol for the indicated time. β-Actin was used as the internal control. (**B**) Real-time analysis of cell death in WT, *Zbp1^−/−^*, and *Zbp1*^ΔZα2^ BMDMs stimulated with ethanol. (**C**) Real-time analysis of cell death in WT and *Zbp1^−/−^* primary hepatocytes stimulated with ethanol. (**D**) Immunoblot analysis of pro- (P45) and activated (P20) CASP1; pro- (P53) and inactivated (P20) GSDMD; pro- (P53) and activated (P34) GSDME; pro- (P55) and cleaved (P18) CASP8; pro- (P35) and cleaved (P17) CASP3; pro- (P35) and cleaved (P20) CASP7; pMLKL, and tMLKL in WT, *Zbp1^−/−^*, and *Zbp1*^ΔZα2^ BMDMs left unstimulated in media (control; Ctrl) or stimulated with ethanol for 4 h. β-Actin was used as the internal control. (**E**) Immunoblot analysis of HMGB1 and LDH in supernatant from WT, *Zbp1^−/−^*, and *Zbp1*^ΔZα2^ BMDMs stimulated with ethanol for the indicated time. β-Actin was used as the loading control. (**F**) Dot blot analysis of Z-form nucleic acid and total DNA (Z-NA and tDNA, respectively) from WT BMDMs left unstimulated in media (Ctrl) or 3 h post-ethanol treatment. (**G**) Representative immunofluorescence micrographs of WT BMDMs treated with ethanol for 3 h and stained for Z-NA (green) and DAPI (nucleus, blue). (**H**) Dot blot analysis of double-stranded and total DNA (dsDNA and tDNA, respectively) from WT BMDMs left unstimulated in media (Ctrl) or 3 h post-ethanol treatment. (**I**) Dot blot analysis of Z-NA and dsDNA and immunoblot analysis of ZBP1 from IgG or anti-ZBP1 immunoprecipitates and from total cell lysates treated with ethanol for 3 h. The data are representative of at least three independent experiments. The data are represented as the mean ± SEM in (**B and C**). The analysis was performed using the two-way ANOVA (or mixed model) in (**B and C**). \*\*\*\**P* < 0.0001. Eth indicates ethanol, and Ctrl indicates control.

Given the elevated ZBP1 expression we observed (**Figure 1A and S1**) and the activation of ethanol-induced PANoptosis molecules (**Figure 2A**), we assessed the involvement of ZBP1, as well as other upstream innate immune sensors and regulators, in ethanol-induced cell death. We found that most innate immune sensors were dispensable for ethanol-induced cell death, including the NOD-like receptors (NLRs) NLRP1, NLRP3, NLRP6, NLRP9b, NLRP12, NLRC1, NLRC2, NLRC3, NLRC4, NLRC5, and NLRX1 (**Figure S4**); Pyrin (*Mefv*), the RNA sensors RIG-I, TLR3, TLR7, and TLR9 (**Figure S5**); and the DNA sensors and signaling molecules cGAS, STING, and AIM2 (**Figure S6**). In contrast, ZBP1 deletion abrogated ethanol-induced cell death in both BMDMs and primary hepatocytes (**Figure 2B and 2C**). Mechanistically, the loss of ZBP1 prevented the cleavage of GSDME, caspases-8, -3, and -7, as well as the phosphorylation of MLKL, thereby limiting the release of DAMPs, including LDH and HMGB1 (**Figure 2D, 2E, and S7A**). These findings suggest that ZBP1 is a central driver of ethanol-induced cell death.

ZBP1 activation is known to be dependent on its sensing ability through its N-terminal Zα2 domain.^28^ We observed that BMDMs lacking the Zα2 domain (*Zbp1*^ΔZα2^) were significantly protected from ethanol-induced cell death, the activation of PANoptosis signaling, and DAMP release (**Figure 2B, 2D, and 2E**). These findings suggest that ZBP1 sensing of nucleic acids plays a central role in the activation of ethanol-induced cell death. To elucidate the mechanism of ZBP1 activation under ethanol exposure, we isolated DNA from ethanol-treated cells and analyzed it using dot blots and immunofluorescence with a Z22 antibody, which selectively binds to Z-NAs. Indeed, Z-NA levels were enriched in response to ethanol treatment (**Figure 2F and 2G**). We also observed enhanced accumulation of dsDNA in response to ethanol treatment (**Figure 2H**). Furthermore, immunoprecipitation experiments showed interactions between ZBP1 and Z-NA, but not dsDNA, in response to ethanol exposure (**Figure 2I**). Collectively, these findings suggest that ethanol-induced Z-NA accumulation contributes to ZBP1-dependent inflammatory cell death.

### IRF9 regulates basal ZBP1 expression to control ethanol-induced cell death

IFN signaling, a critical component of the host immune system, is known to regulate the expression of ZBP1.^29^ To determine the molecular mechanisms involved in IFN signaling-mediated regulation of ZBP1 in ethanol-induced cell death, we sought to identify which IFN regulatory factors (IRFs) were essential. Deletion of IRF9 significantly protected both BMDMs and primary hepatocytes from ethanol-induced death (**Figure 3A–3D**); similarly, deletion of IRF1, a well-known transcriptional regulator of ZBP1,^30^ also reduced cell death (**Figure S8A**). However, the other IRFs tested had no impact on ethanol-induced cell death (**Figure S8B and 8C**). We also conducted RNA-seq analysis on WT and *Irf9*^−/−^ BMDMs and observed a significant decrease in the expression of *Zbp1* in *Irf9*^−/−^ BMDMs compared to WT BMDMs in both untreated and ethanol-treated conditions (**Figure 3E**). RT-PCR and immunoblotting further confirmed that naïve levels of ZBP1 expression were reduced in *Irf9*^−/−^ BMDMs (**Figure 3F and 3G**). Additionally, silencing *Irf9* expression with siRNA reduced basal expression of ZBP1 (**Figure 3H**). Similarly, in overexpression systems, ectopic expression of IRF9 increased the expression of ZBP1 (**Figure 3I**). Together, these data suggest that IRF9, in addition to IRF1, is a key regulator of ZBP1 expression. Furthermore, IRF9 expression was upregulated in patients with both AS and AC (**Figure 3J**), which mirrored the ZBP1 expression data (**Figure 1B and 1C**). Overall, our data suggest that, in addition to IRF1’s role in transcriptional regulation of ZBP1,^30^ IRF9 plays a key role in maintaining baseline ZBP1 expression, enabling ZBP1 to sense ethanol-induced cellular damage and activate inflammatory cell death, PANoptosis.

**Figure 3.**
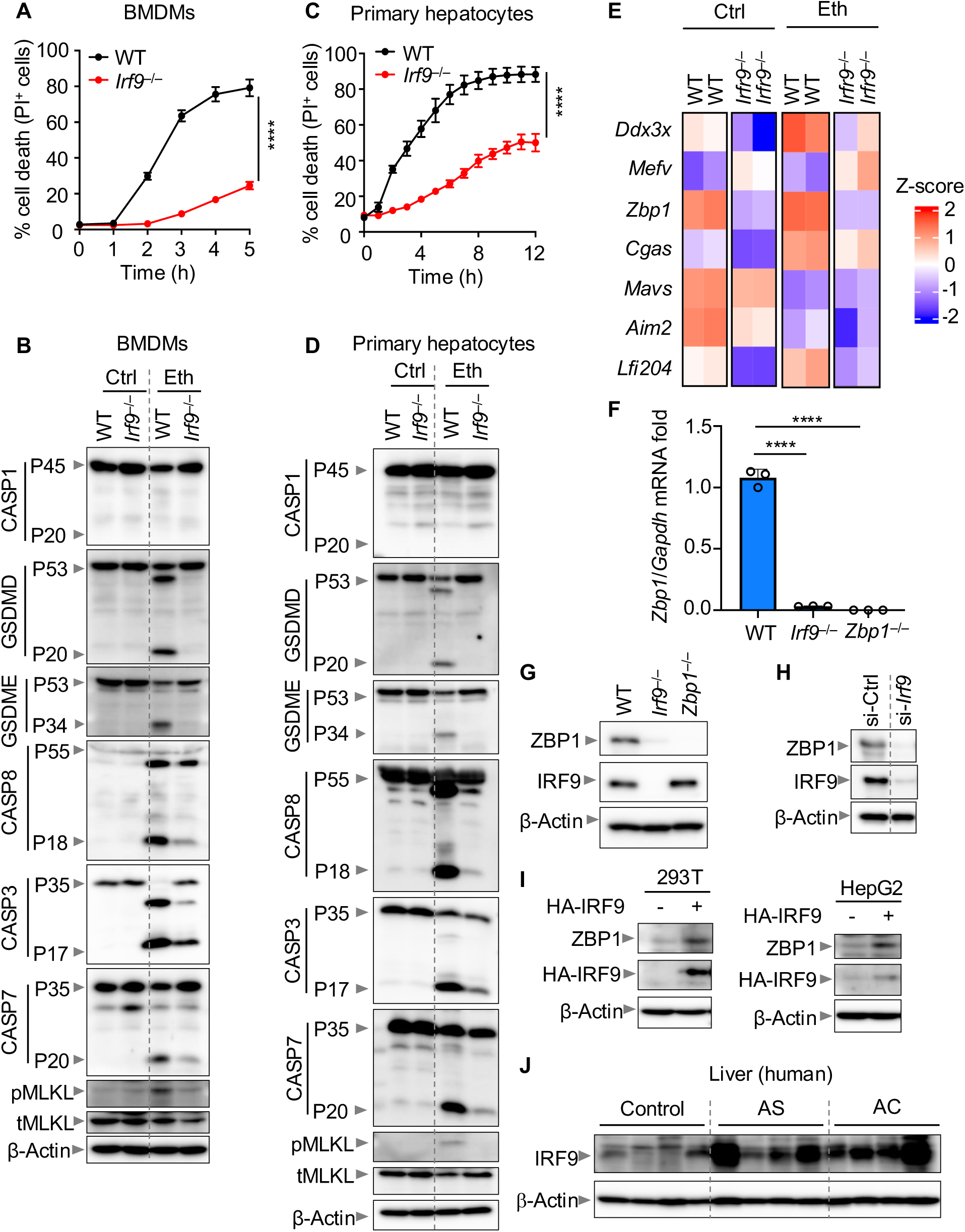
IRF9 regulates basal ZBP1 expression to control ethanol-induced cell death. (**A**) Real-time cell death analysis in wild type (WT) and *Irf9*^−/−^ bone marrow-derived macrophages (BMDMs) stimulated with ethanol. **(B**) Immunoblot analysis of pro- (P45) and activated (P20) caspase-1 (CASP1); pro- (P53) and inactivated (P20) gasdermin D (GSDMD); pro- (P53) and activated (P34) gasdermin E (GSDME); pro- (P55) and cleaved (P18) caspase-8 (CASP8); pro- (P35) and cleaved (P17) caspase-3 (CASP3); pro- (P35) and cleaved (P20) caspase-7 (CASP7); phosphorylated mixed lineage kinase domain-like pseudokinase (pMLKL), and total MLKL (tMLKL) in WT and *Irf9*^−/−^ BMDMs left unstimulated in media (control; Ctrl) or stimulated with ethanol for 4 h. β-Actin was used as the internal control. (**C**) Real-time cell death analysis in WT and *Irf9*^−/−^ primary hepatocytes stimulated with ethanol. **(D**) Immunoblot analysis of pro- (P45) and activated (P20) CASP1; pro- (P53) and inactivated (P20) GSDMD; pro- (P53) and activated (P34) GSDME; pro- (P55) and cleaved (P18) CASP8; pro- (P35) and cleaved (P17) CASP3; pro- (P35) and cleaved (P20) CASP7; pMLKL, and tMLKL in WT and *Irf9*^−/−^ primary hepatocytes left unstimulated in media (control; Ctrl) or stimulated with ethanol for 12 h. β-Actin was used as the internal control. (**E**) Heatmap showing the expression profile of nucleic acid sensors differentially expressed in WT and *Irf9*^−/−^ BMDMs treated with or without ethanol. Blue indicates downregulated genes and red indicates upregulated genes. (**F**) Transcript levels of *Zbp1* in untreated WT, *Irf9*^−/−^, and *Zbp1*^−/−^ BMDMs. (**G**) Immunoblot analysis of ZBP1 and IRF9 in untreated WT, *Zbp1*^−/−^, and *Irf9*^−/−^ BMDMs. β-Actin was used as the internal control. (**H**) Immunoblot analysis of ZBP1 and IRF9 in WT BMDMs treated with control siRNA (si-Ctrl) or *Irf9* siRNA (si-*Irf9*). β-Actin was used as the internal control. (**I**) Immunoblot analysis of ZBP1 and HA-tagged IRF9 (HA-IRF9) in 293T cells and HepG2 cells with or without overexpressed HA-IRF9. β-Actin was used as the internal control. (**J**) Immunoblot analysis of IRF9 in liver tissues from individuals without alcohol-associated liver disease (control), patients with AS, and patients with AC. β-Actin was used as the internal control. The data are representative of at least three independent experiments in (**A–D and F–I**). The data are represented as the mean ± SEM in (**A, C, and F)**. The analysis was performed using the two-way ANOVA (or mixed model) in (**A and C**) or the unpaired t test (one-way ANOVA) in (**F**). \*\*\*\**P* < 0.0001. Eth indicates ethanol, and Ctrl indicates control.

### ZBP1 drives NINJ1, but not the executioners GSDMD, GSDME, or MLKL, to mediate ethanol-induced cell death

Gasdermins and MLKL function as executioners of cell death, resulting in the formation of plasma membrane pores that induce the release of cytosolic content.^31^ NINJ1 then acts as the final gatekeeper, mediating plasma membrane rupture and the release of larger DAMPs.^32^ Based on our observation that multiple executioners, including GSDME and MLKL, were activated in response to ethanol treatment (**Figure 2A**), we sought to determine whether these molecules were required for ethanol-induced cell death. However, we found that *Gsdmd*^−/−^, *Gsdme*^−/−^, and *Mlkl*^−/−^ BMDMs were not protected from ethanol-induced cell death, and the combined deletion of these cell death executioners also failed to provide protection (**Figure S9A–9F**), suggesting that these traditional lytic executioners are not required for mediating ethanol-induced cell death. NINJ1 can also be required for the execution of cell death under various conditions, including exposure to heat shock and cumene hydroperoxide.^32–34^ Therefore, we sought to determine whether NINJ1 was critical for ethanol-induced cell death. Deletion of NINJ1 provided significant protection from lytic cell death in response to ethanol treatment (**Figure 4A and 4B**), and comparable results were observed using *Ninj1* siRNA-based silencing (**Figure 4C**). Additionally, blocking NINJ1 clustering with glycine^35^ prevented ethanol-induced cell death (**Figure 4D**) and the release of inflammatory DAMPs (**Figure 4E**). Furthermore, glycine treatment protected *Gsdmd*^−/−^*Gsdme*^−/−^*Mlkl*^−/−^ BMDMs from cell death (**Figure 4F**), suggesting that NINJ1, rather than GSDMD, GSDME, or MLKL, was the downstream molecule inducing cell death in response to ethanol. Moreover, we observed NINJ1 oligomerization in response to ethanol treatment, which was reduced in *Zbp1^−/−^* BMDMs (**Figure 4G**), consistent with the reduced cell death observed in these cells (**Figure 2B**). Overall, these findings suggest that ethanol promotes NINJ1 oligomerization that is regulated by ZBP1 and leads to NINJ1-dependent plasma membrane rupture.

**Figure 4.**
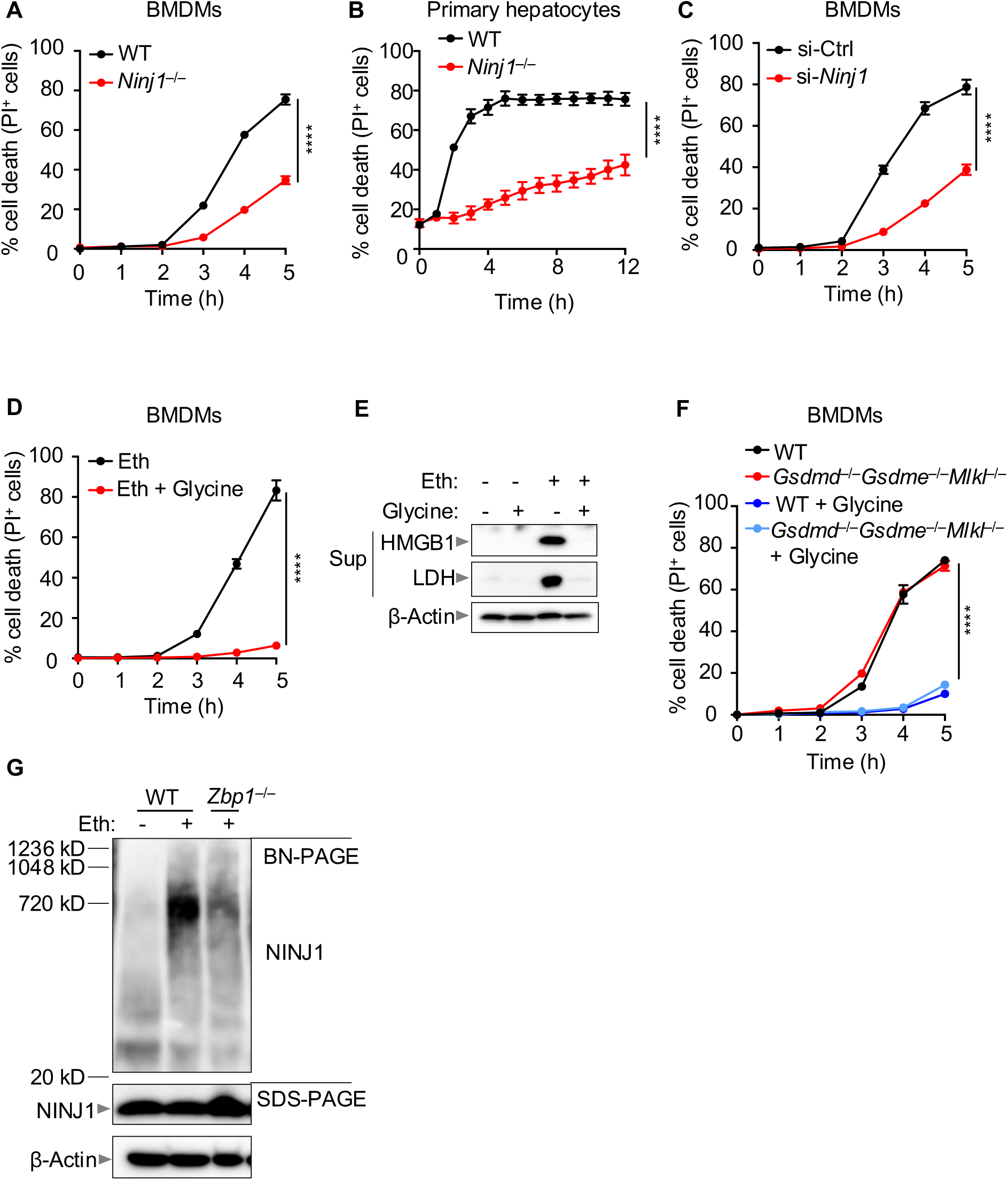
ZBP1 drives NINJ1, but not the executioners GSDMD, GSDME, or MLKL, to mediate ethanol-induced cell death. (**A**) Real-time cell death analysis in wild type (WT) and *Ninj1*^−/−^ bone marrow-derived macrophages (BMDMs) stimulated with ethanol. (**B**) Real-time cell death analysis in WT and *Ninj1*^−/−^ primary hepatocytes stimulated with ethanol. (**C, D**) Real-time analysis of cell death in (**C**) control siRNA (si-Ctrl)- and *Ninj1* siRNA (si-*Ninj1*)-treated, and (**D**) glycine-treated BMDMs stimulated with ethanol. (**E**) Immunoblot analysis of HMGB1 and LDH in the supernatant (Sup) from BMDMs treated with or without glycine and ethanol for 4 h. (**F**) Real-time analysis of cell death in WT and *Gsdmd*^−/−^*Gsdme*^−/−^*Mlkl*^−/−^ BMDMs stimulated with ethanol and treated with or without glycine. (**G**) Blue native polyacrylamide gel electrophoresis (BN-PAGE) and SDS-PAGE analysis of NINJ1 in WT and *Zbp1^−/−^* BMDMs stimulated with ethanol for 3 h. β-Actin was used as the internal control. The data are representative of at least three independent experiments. The data are represented as the mean ± SEM in (**A–D and F**). The analysis was performed using the two-way ANOVA (or mixed model) in (**A–D and F**). **** *P* < 0.0001. Eth indicates ethanol.

### ZBP1 drives pathogenesis in chronic and acute ethanol-induced liver disease in mice

Given the significance of the ZBP1 signaling pathway in driving ethanol-induced cell death, we next sought to determine whether this pathway contributed to pathology in a murine chronic-plus-binge ethanol diet model (**Figure 5A**). We found that WT mice on an ethanol diet developed hepatic macrovesicular steatosis, which was characterized by lipid accumulation in hepatocytes as vacuoles (**Figure 5B and 5C**). This observation was consistent with elevated serum levels of alanine aminotransferase (ALT) and aspartate aminotransferase (AST) following ethanol administration compared to the control group (**Figure 5D and 5E**). In contrast, *Zbp1^−/−^*mice exhibited milder ethanol-associated liver damage, characterized by reduced vacuole accumulation and lower serum ALT and AST levels (**Figure 5B–E and S10A**). Immunohistochemistry further confirmed that *Zbp1^−/−^*mice had reduced cleaved caspase-3 in their liver tissues compared with WT mice (**Figure 5F**), suggesting reduced cell death was occurring in *Zbp1^−/−^* mice response to the ethanol diet. These findings suggest that ZBP1 plays a critical role in the progression of ALD in vivo.

**Figure 5.**
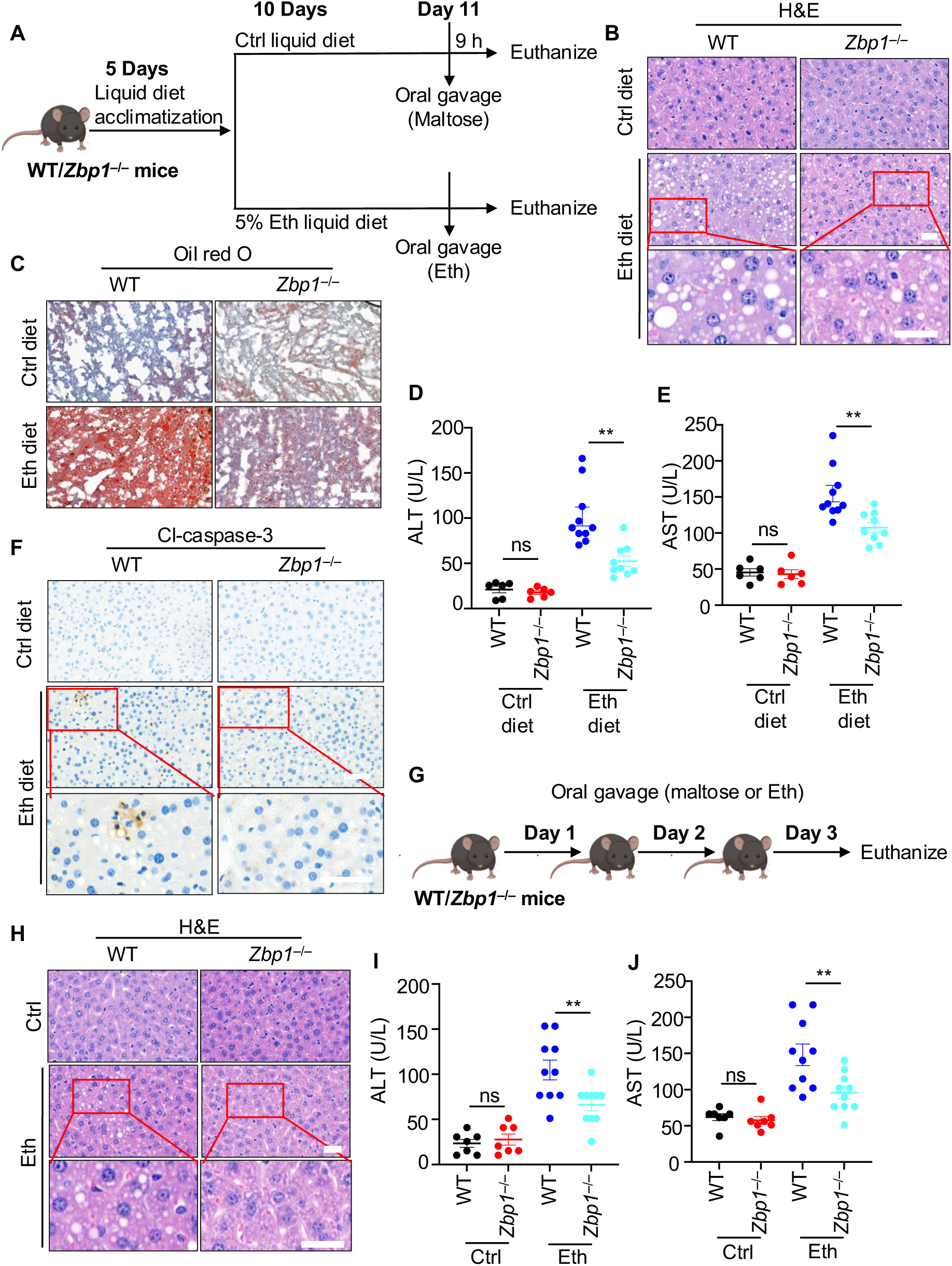
ZBP1 drives pathogenesis in chronic and acute ethanol-induced liver disease in mice. (**A**) Schematic illustrating the chronic-plus-binge ethanol diet-induced liver disease model. (**B**) Hematoxylin and eosin (H&E)-stained liver sections from control diet (Ctrl diet)- and ethanol diet (Eth diet)-fed groups of wild type (WT) and *Zbp1*^−/−^ mice. Red insets represent the enlarged images. Scale bars, 25 µm. (**C**) Oil red O staining of liver sections from Ctrl diet- and Eth diet-fed groups of WT and *Zbp1*^−/−^ mice. Scale bar, 100 µm. (**D and E**) Comparison of serum alanine transaminase (ALT) and aspartate transaminase (AST) levels between WT and *Zbp1*^−/−^ mice in the Ctrl diet- and Eth diet-fed groups. (**F**) Immunohistochemistry staining of cleaved-caspase-3 (Cl-caspase-3) in liver sections from Ctrl diet- and Eth diet-fed groups of WT and *Zbp1*^−/−^ mice. Red insets represent the enlarged images. Scale bars, 50 µm. (**G**) Schematic illustrating the experimental animal model of acute ethanol-induced liver disease. (**H**) H&E-stained liver sections from WT and *Zbp1*^−/−^ mice in the Ctrl and Eth-treated groups. Red insets represent the enlarged images. Scale bars, 25 µm. (**I and J**) Comparison of serum ALT and AST levels between WT and *Zbp1*^−/−^ mice in the Ctrl and Eth-treated groups. The data are representative of two independent experiments. The data are represented as the mean ± SEM in (**D, E, I, and J**). The analysis was performed using the unpaired t test (one-way ANOVA) in (**D, E, I, and J**). ns, not significant; \*\**P* < 0.01.

To further evaluate the role of ZBP1 in the development of ALD, we used a murine model of acute ethanol-induced liver disease (**Figure 5G**). Consistent with the results observed in the chronic-plus-binge ethanol diet model, we observed that *Zbp1^−/−^* mice were significantly protected from ethanol-associated liver injury. This was evident from the reduced vacuole formation in the liver, and significantly reduced serum ALT and AST levels as compared to WT mice (**Figure 5H–J and S10B**). Taken together, these findings suggest that ZBP1 is critical for both the early sensing of ethanol-induced damage and the pathogenesis of chronic and acute ALD.

### NINJ1 serves as a biomarker for alcohol-associated liver disease

Our observations suggest that the ZBP1-mediated pathway contributed to the progression of ALD in both human patients and murine ALD models. Given the critical role we identified for NINJ1 in mediating cell death in response to ethanol, we investigated NINJ1’s role in ALD. Analysis of publicly available datasets identified that NINJ1 expression was increased in patients with AH compared to healthy controls across multiple datasets (GSE28619 and GSE155907^36,37^) (**Figure 6A**). Furthermore, liver homogenates from the murine model of ALD and patients with AS and AC demonstrated that NINJ1 protein levels were also increased during disease (**Figure 6B and 6C**). Consistent with our observations in human liver tissues, we also found that NINJ1 expression was elevated in the serum samples of patients with both AS and AC (**Figure 6D**). This mirrored the expression pattern of ZBP1 in patients’ serum samples (**Figure 6E and S11A**). These results show that ZBP1 and NINJ1 expression was moderately elevated in patients with AS, with ZBP1 being elevated further in patients with AC. Collectively, our findings suggest that NINJ1, in combination with ZBP1, shows potential as a circulating biomarker for ALD diagnosis and as a promising therapeutic target for ALD intervention.

**Figure 6.**
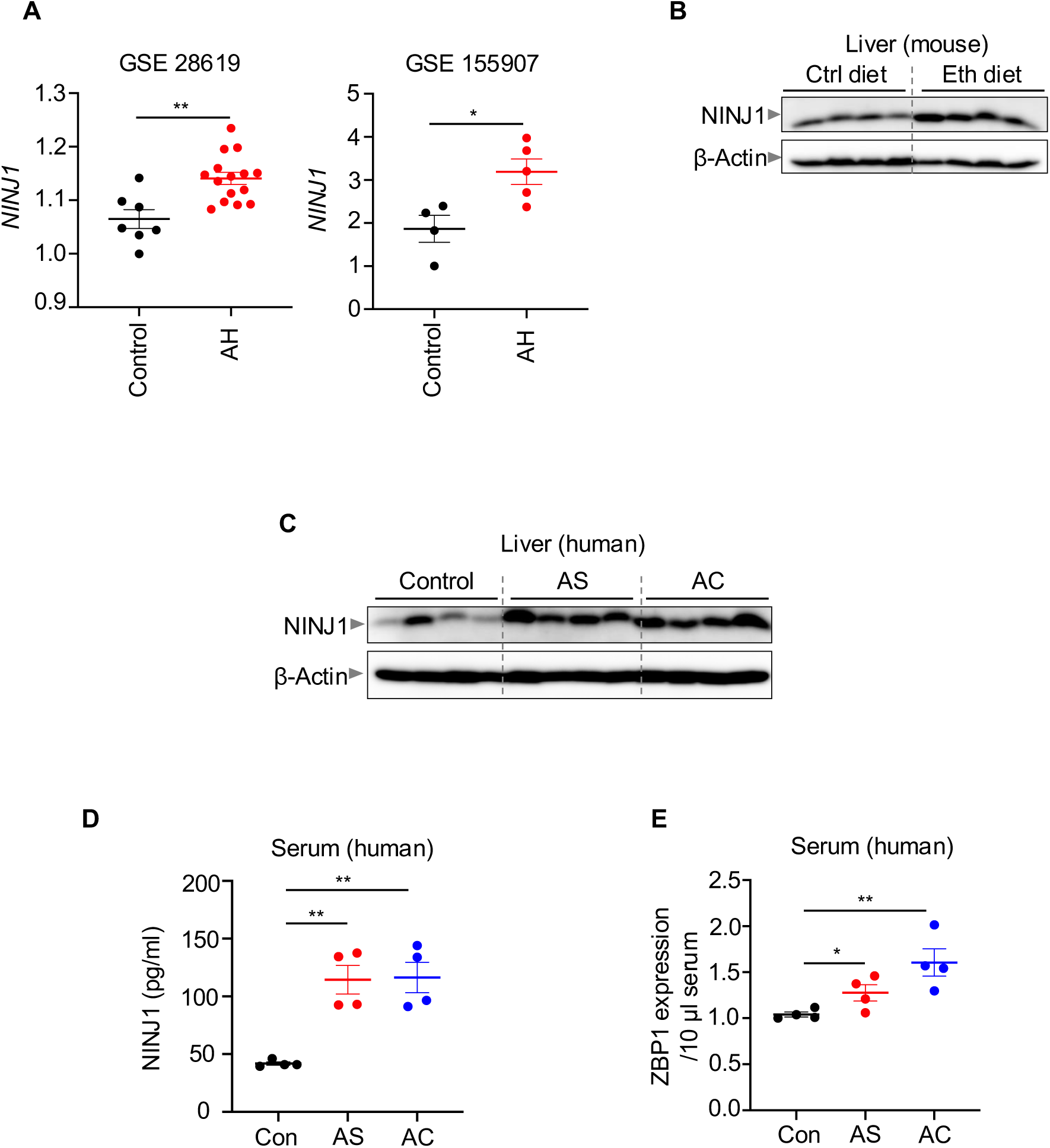
NINJ1 serves as a biomarker for alcohol-associated liver disease. (**A**) Relative expression of *NINJ1* in liver tissues from individuals without alcohol-associated liver disease (control) and patients with alcoholic hepatitis (AH), analyzed using publicly available datasets (GSE28619 and GSE155907). (**B**) Immunoblot analysis of NINJ1 expression in the liver tissues from wild type (WT) control (Ctrl diet) and ethanol-administered (Eth diet) groups in the murine chronic-plus-binge ethanol diet model. β-Actin was used as the internal control. (**C**) Immunoblot analysis of NINJ1 expression in the liver tissues of individuals without alcohol-associated liver disease (control) and patients with alcohol-associated steatosis (AS) or alcoholic cirrhosis (AC). (**D**) ELISA analysis of NINJ1 expression in the serum samples from individuals without alcohol-associated liver disease (control) and patients with AS or AC. (**E**) Densitometry analysis of ZBP1 expression in the serum samples of individuals without alcohol-associated liver disease (control) and patients with AS or AC. The analysis was performed using the unpaired t test (one-way ANOVA) in (**A, D, and E**). \**P* < 0.05 and \*\**P* < 0.01.

## Discussion

Activation of the innate immune response and cell death pathways is a critical component of host defense to eliminate pathogens. However, excess activation in response to sterile triggers, such as ethanol, can lead to inflammation and tissue damage.^3–7^ Therefore, pathways that drive inflammation are associated with pathology and disease, including in ALD. With nearly two billion individuals consuming alcohol worldwide, alcohol remains a leading cause of disease and a major risk factor for premature mortality among individuals aged 15 to 49.^38^ Furthermore, 75 million individuals experience alcohol-associated disorders, including ALD,^2^ which accounts for a significant proportion of global alcohol-associated mortality.^2,39,40^ Despite ongoing clinical investigations, there are currently no FDA-approved treatments for ALD, making it critical to elucidate the molecular mechanisms of ethanol-induced inflammation and tissue injury to identify therapeutic targets.

Our results suggest that ethanol induced robust inflammatory cell death in both immune and non-immune cells, including liver hepatocytes and Kupffer cells. We found that the innate immune nucleic acid sensor ZBP1 played a central role in the regulation of ethanol-induced cell death and ALD. ZBP1 was initially characterized as the innate immune sensor for influenza A virus (IAV),^16^ and it is now known to be a key regulator of PANoptosis that can drive inflammation in response to IAV infection,^16^ IFN therapy during β-coronavirus infection,^29^ and the combination of NEIs and IFN.^17^ Furthermore, together with cGAS, ZBP1 can sense mitochondrial DNA to promote cardiotoxicity.^41^ However, the role of ZBP1 in the onset and progression of ALD was previously unclear. Additionally, while ZBP1 expression is most well known to be regulated by IRF1,^42^ IRF9-mediated ZBP1 expression may be a key regulatory step in ALD, where we found that IRF9 expression was upregulated. Given that ZBP1 expression was consistently elevated in the liver tissue from different stages of ALD, and that its expression increased with disease progression, ZBP1 may be a useful biomarker for ALD and disease progression. Furthermore, ZBP1-deficient mice were protected from many of the clinical features of ALD in response to both chronic and acute ethanol consumption, highlighting ZBP1 as a promising therapeutic target for ALD. Our findings also indicate that ethanol exposure led to Z-NA accumulation, and this accumulation was detected by ZBP1’s Zα2 domain, ultimately triggering ethanol-induced cell death, PANoptosis.

Downstream of ZBP1 sensing, our results indicate a distinct role for NINJ1 in regulating ethanol-induced cell death, rather than the conventional lytic cell death executioners GSDMD, GSDME, or MLKL. NINJ1 can be activated downstream of caspase-8,^33^ and we observed robust ZBP1-dependent caspase-8 activation in response to ethanol. NINJ1 oligomerization is required for membrane lysis,^35^ and our results suggest that preventing this oligomerization blocked cell death and DAMP release. These DAMPs are likely contributors to the progression of inflammation and tissue damage in ALD. Moreover, our analyses indicated that NINJ1 expression was upregulated in liver tissues and serum samples from patients with AS and AC, further implicating a central role for NINJ1-mediated cell membrane rupture in driving inflammation and ALD.

Overall, our study provides a comprehensive characterization of the molecular mechanisms underlying inflammatory cell death in response to ethanol. Our findings highlight the critical roles of ZBP1 and NINJ1 in the development of ALD. Furthermore, NINJ1, along with ZBP1, may serve as promising circulating biomarkers for ALD diagnosis and progression monitoring, as well as potential therapeutic targets to limit inflammation and pathology across the disease spectrum.

## ACKNOWLEDGMENTS

The human liver specimens used in this study were obtained from the University of Kansas Liver Tissue Biorepository, supported by grant P20GM144269 from the National Institute of General Medical Sciences. We acknowledge the patients who generously donated valuable specimens for research, as well as the physicians, nurses, and researchers who procured the specimens. The *Irf9*^−/−^ mutant mouse was provided by RIKEN BRC through the National Bio-Resource Project of the MEXT, Japan. Previously published datasets that were analyzed for this study are deposited in the Gene Expression Omnibus (GEO) database (accession IDs: GSE143318, GSE28619, and GSE155907). We thank all the members of the Kanneganti laboratory for their scientific input, comments, and suggestions, and Anu Sharma, PhD, and Sara Resende, PhD, for scientific editing support. This research was supported by NIH grants AI101935, AI124346, AI160179, AR056296, and CA253095, and support from the American Lebanese Syrian Associated Charities to Dr. Kanneganti. The content is solely the responsibility of the authors and does not necessarily represent the official views of the National Institutes of Health.

## METHODS

### Mice

*Nlrp3*^−/−^,^43^ *Aim2*^−/−^,^44^ *Nlrp1*^−/−^,^45^ *Nlrc4*^−/−^,^46^ *Mefv*^−/−^,^47^ *Nlrc1*^−/−^,^48^ *Nlrc2*^−/−^,^49^ *Nlrc3*^−/−^,^50^ *Nlrc5*^−/−^,^51^ *Nlrx1*^−/−^,^52^ *Nlrp6*^−/−^,^53^ *Nlrp9b*^−/−^,^54,55^ *Nlrp12*^−/−^,^56^ *Rigi*^−/−^,^57^ *Tlr3*^−/−^,^58^ *Tlr7*^−/−^,^59^ *Tlr9*^−/−^,^60^ *Cgas*^−/−^,^61^ *Sting*^−/−^,^62^ *Irf1^-/-^*,^63^ *Irf8*^−/−^,^64^ *Irf3*^−/−^,^65^ *Irf5*^−/−^,^66^ *Irf7*^−/−^,^67^ *Irf3/7*^−/−^,^68^ *Irf9*^−/−^,^69^ *Gsdmd*^−/−^,^68^ *Gsdme*^−/−^,^54^ *Mlkl*^−/−^,^70^ *Gsdmd*^−/−^*Gsdme*^−/−^,^17^ *Gsdmd*^−/−^*Mlkl*^−/−^,^20^ *Gsdmd*^−/−^*Gsdme*^−/−^*Mlkl*^−/−^,^63^ *Ninj1*^−/−^,^33^ *Zbp1*^−/−^,^71^ and *Zbp1*^ΔZα2^ ^72^ mice have been previously described. *Tlr3/7/9*^−/−^ mice were generated at our facility by crossing *Tlr3*^−/−^ and *Tlr7*^−/−^ mice, then by crossing *Tlr3/7*^−/−^ mice with *Tlr9*^−/−^ mice. All mice were generated on or extensively backcrossed to the C57/BL6 background and were bred at the Animal Resources Center at the St. Jude Children’s Research Hospital under specific pathogen-free conditions. Both male and female age- and sex-matched 8- to 10-week-old mice were used in this study. Mice were maintained on a 12 h light-dark cycle and were fed standard chow. Animal studies were conducted under protocols approved by the St. Jude Children’s Research Hospital Committee on the Use and Care of Animals.

### Human liver samples

Frozen liver tissue samples (including alcoholic cirrhosis, alcoholic steatosis/steatohepatitis, and normal control liver samples) were provided by the University of Kansas Liver Tissue Biorepository, supported by grant P20GM144269 from the National Institute of General Medical Sciences. The studies were performed with approval and according to the relevant guidelines established at the institution of collection.

### Cell culture

Primary mouse bone marrow-derived macrophages (BMDMs) were generated from the bone marrow of wild type and indicated mutant mice and grown for 5–6 days in IMDM medium (Thermo Fisher Scientific, 12440-053) containing 1% non-essential amino acids (Thermo Fisher Scientific, 11140-050), 10% heat-inactivated fetal bovine serum (HI-FBS; Biowest, S1620), 30% L929 conditioned media, and 1% penicillin and streptomycin (Thermo Fisher Scientific, 15070-063). BMDMs were then seeded into antibiotic-free media at a density of 1×10^6^ cells/well in 12 well plates or 5×10^5^ cells/well in 24 well plates and incubated overnight before use.

THP-1 cells (ATCC, TIB-202) were cultured in RPMI medium (Corning, 10-040-CV) containing 10% heat inactivated HI-FBS and 1% penicillin and streptomycin and treated with 100 ng/ml phorbol 12-myristate 13-acetate (PMA) overnight to differentiate into macrophages. 293T cells (ATCC, CRL-3216) were cultured in DMEM (Gibco, 11995-065) supplemented with 1% penicillin and streptomycin and 10% heat inactivated HI-FBS. The human HepG2 cells (ATCC, HB-8065) were cultured in Eagle’s Minimum Essential Medium (EMDM) (ATCC, 30-2003) containing 1% penicillin and streptomycin and 10% heat inactivated HI-FBS.

Primary hepatocytes were isolated from mice, then purified and cultured as previously described in Star Protocols.^73^ Briefly, mice were anesthetized and perfused. The perfused livers were removed from the mice, then broken up using forceps, and gently mixed with a cell lifter. The liver digests were filtered through a 70 μm cell strainer (Fisher Scientific, 23363548) and washed with Hepatocyte Wash Medium (Gibco, 17704024). The homogenate was centrifuged at 50 × g for 2 min three times with no acceleration and no brake. Hepatocytes were collected from cell pellets, and cell viability was determined. Then, hepatocytes were counted and plated in HepatoZYME-SFM medium (Gibco, 17705021) for further analysis.

Rat Kupffer cells used in the study were purchased from Thermo Fisher Scientific (RTKCCS) and handled according to the manufacturer’s instructions.

For the overexpression system, 293T cells and HepG2 cells were seeded into 12 well plates and transfected with 1 μg *IRF9* plasmid (Addgene, #11614) and incubated for 24–36 h. Cell lysates were collected and used for analysis.

### Cell stimulation

The following ligands were used to stimulate primary mouse HepG2 cells, THP-1 cells, primary hepatocytes, Kupffer cells, and BMDMs: 5% or the indicated concentration of ethanol (vol/vol; Fisher, 04-355-451) and 5 mM glycine (Sigma-Aldrich, G7126) for the indicated times. A total of 250 mL or 500 mL of media containing the indicated ligands was utilized for stimulation in 24- or 12-well plates, respectively.

### Real-time imaging for cell death

The kinetics of cell death were analyzed using the IncuCyte S3 (Sartorius) live-cell automated system. BMDMs (5×10^5^ cells/well) and THP-1 cells (5×10^5^ cells/well) were seeded in 24-well tissue culture plates and treated with the indicated stimuli. Cells were stained with propidium iodide (PI; Life Technologies, P3566) following the manufacturer’s protocol. The plate was scanned for the indicated durations, and fluorescent and phase-contrast images were acquired in real-time every 1 h. PI-positive dead cells are marked with a red mask for visualization. The image analysis, masking, and quantification of dead cells (PI^+^ cells) were performed using the software package supplied with the IncuCyte imager. In some cases, a single WT was used to compare against multiple knockout lines inoculated and stimulated simultaneously in the same experiment. The data are plotted separately in the figures for optimal visualization; therefore, the same WT cell death curve appears in multiple panels.

### Immunoblot analysis

For probing caspase activation, cell lysates and culture supernatants were combined in 50 μl caspase lysis buffer (containing 1× protease inhibitors, 1× phosphatase inhibitors, 10% NP-40, and 25 mM DTT) and 4× sample loading buffer (containing SDS and 2-mercaptoethanol). For immunoblot analysis of signaling activation, culture supernatants were removed, and cells were washed once with DPBS (Thermo Fisher Scientific, 14190-250), followed by lysis in RIPA buffer and sample loading buffer. For immunoblot analysis of LDH and HMGB1 in the supernatant, the supernatant was collected and centrifuged at 8,000 × g for 2 min, removing cell debris. The obtained supernatant was combined with a 4× sample loading buffer. Proteins were separated by electrophoresis through 8–12% polyacrylamide gels, followed by electrophoretic transfer of proteins onto PVDF membranes (Millipore, IPVH00010). The membranes were blocked by incubation with 5% skim milk then incubated with primary antibodies against: anti-caspase-1 (AdipoGen, AG-20B-0044, 1:1,000), anti-GSDMD (Abcam, ab209845, 1:1,000), anti-GSDME (Abcam, ab215191, 1:1,000), anti-caspase-8 (CST, 4927, 1:1,000), anti-cleaved caspase-8 (CST, 8592, 1:1,000), anti-caspase-3 (CST, 9662, 1:1,000), anti-cleaved caspase-3 (CST, 9661, 1:1,000), anti-caspase-7 (CST, 9492, 1:1,000), anti-cleaved caspase-7 (CST, 9491, 1:1,000), anti-pMLKL (CST, 37333, 1:1,000), anti-MLKL (Abgent, AP14272b, 1:1,000), anti-ZBP1 (AdipoGen, AG-20B-0010, 1:1,000), anti-NINJ1 (ABclonal A16406, 1:1,000), anti-LDHA (Proteintech, 19987-1-AP, 1:1,000), anti-HMGB1 (abcam, ab79823, 1:1,000), anti-HA (Millipore, 05-904, 1:1,000), anti-IRF9 (Proteintech, 14167-1-AP, 1:1,000), and anti-β-actin (Proteintech, 66009-1-IG, 1:5,000). Membranes were then washed and probed with the appropriate horseradish peroxidase (HRP)–conjugated secondary antibodies (Jackson ImmunoResearch Laboratories, anti-rabbit [111-035-047] 1:5,000 and anti-mouse [315-035-047] 1:5,000 for 1 h. Proteins were visualized using Immobilon Forte Western HRP Substrate (Millipore, WBLUF0500) or SuperSignal West Femto Maximum Sensitivity Substrate (Thermo Fisher Scientific, 34096) on the GE Amersham Imager 600. Images were analyzed with ImageJ (v1.53a).

### Gray analysis of protein expression

Densitometry was calculated using the band intensity from the western blots. The quantification of the indicated protein expression was performed using ImageJ v1.54i software. The ZBP1 protein expression was normalized to the β-actin protein expression (loading control) in Figure 1C or sample volume in Figure 6E for each lane.

### Dot blot analysis

DNA extraction was performed according to the manufacturer’s instructions (69506, Qiagen). A dot blot for Z-NA or dsDNA analysis was performed with total DNA isolated from the cells with an anti-Z-NA antibody (Z22; Absolute Ab00783) or an anti-dsDNA antibody (Abcam, Ab27156). Equal volumes (2 μL containing 1 μg of DNA) of the DNA were dotted on a Hybond N+ membrane (GE Healthcare, RPN203B), dried, and autocrosslinked in a UV stratalinker 2400 (Stratagene, 5496A UV) two times. The membrane was then blocked in 5% milk in TBST for 1 h and probed with a Z-NA or dsDNA antibody at 4°C overnight. The membrane was washed and probed with the horseradish peroxidase (HRP)–conjugated secondary antibody (Jackson ImmunoResearch Laboratories, anti-mouse [315-035-047] 1:5,000) for 1 h. The membrane was then processed for immunoblot analysis. The total DNA was detected using 0.2% methylene blue (Sigma, M9140).

### Immunoprecipitation

For immunoprecipitation, 4×10^7^ BMDMs were seeded and stimulated with ethanol for 3 h. The cell lysates were prepared in an ice-cold lysis buffer containing 20 mM Tris-HCl (pH 7.4), 150 mM NaCl, 1% Triton X-100, 10% glycerol, 1 mM Na_3_VO_4_, 2 mM PMSF, EDTA-free protease inhibitor cocktail (Thermo Fisher Scientific, A32965), and phosphatase inhibitor cocktail (Sigma, 4906845001). Immunoprecipitation was performed by incubating the cell lysates with anti-ZBP1 antibody (AdipoGen, AG-20B-0010) or anti-IgG antibody (CST, 68860) and Protein A/G magnetic beads (Thermo Fisher Scientific, #88802,). The samples were then subjected to immunoblot and dot blot analysis.

### Immunofluorescence staining

BMDMs were fixed in 4% paraformaldehyde for 10 min at room temperature, followed by permeabilization in 0.2% Triton X-100 for 10 min. Cells were then blocked in 5% normal goat serum (Life Technologies, 01-6201) for 1 h at room temperature. Samples were incubated with an anti-Z-NA antibody (1:250) overnight at 4°C. Cells were then washed three times with PBS and incubated with AlexaFluor 488-conjugated antibody against mouse IgG (Invitrogen, A11029, 1:1000) and counterstained with DAPI (Biotium, 40043) for 1 h at room temperature. Cells were washed three times with PBS and imaged using a Leica SP8 confocal microscope.

### BN-PAGE

The BMDMs were lysed with native-PAGE lysis buffer (1% Digitonin, 150 mM NaCl, 50 mM Tris pH 7.5, and 1× complete protease inhibitor). After centrifuging at 20,800 × g for 30 min, lysates were mixed with 4× NativePAGE sample buffer (Thermo Fisher Scientific, BN2003) and Coomassie G-250 (Thermo Fisher Scientific, BN2008). Then, samples were subjected to BN-PAGE using NativePAGE 3–12% Gel (Thermo Fisher Scientific, BN1001BOX).

### siRNA-mediated gene silencing

A total of 5 nmol of siRNA (si-*Irf9* (Horizon Discovery, M-062207-00-0005); si-*Ninj1* (Horizon Discovery, M-048149-01-0005); si-Ctrl (Horizon Discovery, D-001210-01-05)) was dissolved in sterile nuclease-free water to a final concentration of 50 μM, and 0.5 μl of siRNA was added to 1×10^6^ BMDMs. Electroporation was performed using the neon transfection system (Invitrogen), with parameters of 1500 V, 1 pulse, and 20 ms width. After electroporation, BMDMs were immediately transferred into 24-well plates with a seeding density of 5×10^5^ cells per well. Two days after transfection, BMDMs were stimulated as required to assess cell death.

### RNAseq analysis

The mouse transcripts were analyzed by profiling BMDMs derived from WT and *Irf9^−/−^* mice. Briefly, the RNA was extracted from WT and *Irf9^−/−^* BMDMs using the RNeasy Mini Kit (QIAGEN, 74014) according to the manufacturer’s instructions. RNA sequencing data were processed by the St. Jude Center for Applied Bioinformatics AutoMapper pipeline. The raw FASTQ data were quality filtered, and the adapter sequences were removed using TrimGalore v0.6.3 (https://github.com/FelixKrueger/TrimGalore). STAR^74^ v2.7.9a was then used to align the human reference genome (GRCh38) in a splice-site aware manner using Gencode v31 primary assembly annotations. Gene and isoform level quantifications were computed using RSEM^75^ v1.3.3.

The role of nucleic acid sensors in ALD was assessed using data from the Gene Expression Omnibus (GEO) database (accession ID: GSE143318). GSE143318 comprises gene expression profiles of liver tissues from 13 patients with alcoholic hepatitis, 5 with alcoholic cirrhosis, and 7 control donors. The NINJ1 expression profile was obtained from the GEO database (accession IDs: GSE28619 and GSE155907). The dataset GSE28619 included 7 control subjects and 15 patients with alcoholic hepatitis. The dataset GSE155907 included 4 control subjects and 5 patients with alcoholic hepatitis.

The analysis included quality control steps, including normalized quantiles using the ‘normalize. quantiles’ function from the preprocess Core v1.58.0 package when the counts were not normalized, followed by log2 transformation for downstream differential expression analysis. The differential expression analysis was performed using the limma v3.52.1 package in R v4.1.1.

### RT-PCR analysis

Total RNA was extracted using TRIzol (Thermo Fisher Scientific, 15596026). cDNA was synthesized with 500 ng of extracted RNA using the High-Capacity cDNA Reverse Transcription Kit (4368814, Applied Biosystems). Real-time quantitative PCR was performed on an Applied Biosystems 7500 real-time PCR instrument using SYBR Green (4368706, Applied Biosystems). The primer sequences are as follows: mZBP1-Forward primer: CTCCTGCAATCCCTGAGAACT; Reverse primer: GGCTACATGGCAAGACTATGTC; mGAPDH-Forward primer: CGTCCCGTAGACAAAATGGT; Reverse primer: TTGATGGCAACAATCTCCAC.

### Chronic-plus-binge ethanol diet model

Age- and sex-matched, 8- to 10-week-old WT and *Zbp1*^−/−^ mice were cohoused and initially provided with unrestricted access to the control Lieber-DeCarli diet ad libitum for 5 days to acclimatize them to a liquid diet. Then, ethanol-fed groups were fed with an ethanol Lieber-DeCarli diet containing 5% (vol/vol) ethanol, while the control groups were pair-fed a control diet that isocalorically substituted maltose dextrin for ethanol for 10 days. On day 11, pair-fed mice were gavaged with 5 g/kg maltose, while ethanol-fed mice were gavaged with 5 g/kg ethanol. Mice were euthanized 9 h later. Blood samples were collected, and liver samples were harvested and stored appropriately for further analysis.

### Acute binge model

Age- and sex-matched 8- to 10-week-old WT and *Zbp1*^−/−^ mice were cohoused and administered a daily oral gavage of ethanol at 5 g/kg body weight (20% vol/vol in sterile phosphate buffer saline) or equal calorie maltose for three days. Mice were euthanized 9 h after the final alcohol gavage. Blood samples were collected, and liver samples were harvested and stored appropriately for further analysis.

### Histopathology

Murine liver tissues from the in vivo ALD models were fixed in 10% formalin, processed, and embedded in paraffin according to standard procedures. Tissue sections (5 mM) were stained with hematoxylin and eosin (H&E) or with cleaved-caspase-3 antibody (CST, #9664). For Oil-Red O staining, liver tissues were embedded in Tissue-Tek O.C.T. Compound (SAKURA 4583), sectioned, and subjected to Oil-Red O staining. The histological results were examined by a pathologist blinded to the experimental groups.

### Enzyme-linked immunosorbent assay (ELISA)

Patient serum was assessed using ELISA kits according to the manufacturer’s instructions (biorbyt, orb404939).

### Statistical analysis

GraphPad Prism 9.0 software was used for data analysis. Data are presented as mean ± SEM. The student’s t tests (two-tailed) or two-way ANOVA (or mixed model) were used to determine the statistical significance. *P* values less than 0.05 were considered statistically significant, where \**P* < 0.05, \*\**P* < 0.01, \*\*\**P* < 0.001, and \*\*\*\**P* < 0.0001. At least two independent experiments were performed to generate each dataset. The statistical test employed for each experiment is indicated in the figure legends.

