## Supplemental Figures and Legends for "The critical role of the ZBP1-NINJ1 axis and IRF1/IRF9 in ethanol-induced cell death, PANoptosis, and alcohol-associated liver disease"

**A**

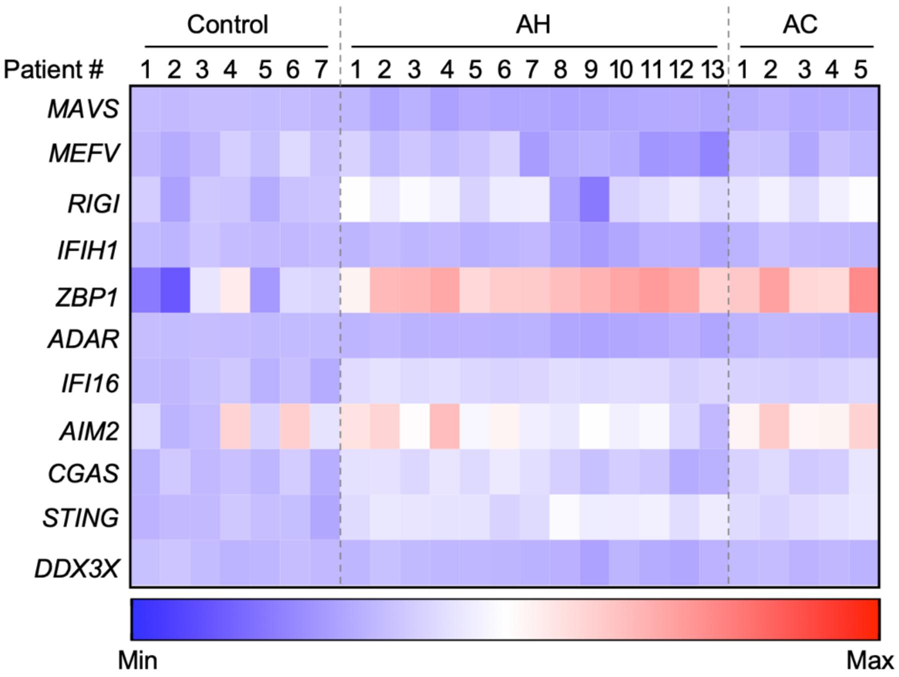


**Figure S1. ZBP1 expression is upregulated across individual patients with alcohol-associated liver disease**

(**A**) Heatmap showing the expression profile of nucleic acid sensors in the liver tissues from individuals without alcohol-associated liver disease (control), patients with alcoholic hepatitis (AH), and patients with alcoholic cirrhosis (AC). The heatmap represents differentially expressed genes (fold change), where downregulated genes are in blue and upregulated genes in red, compared with the average expression in controls.

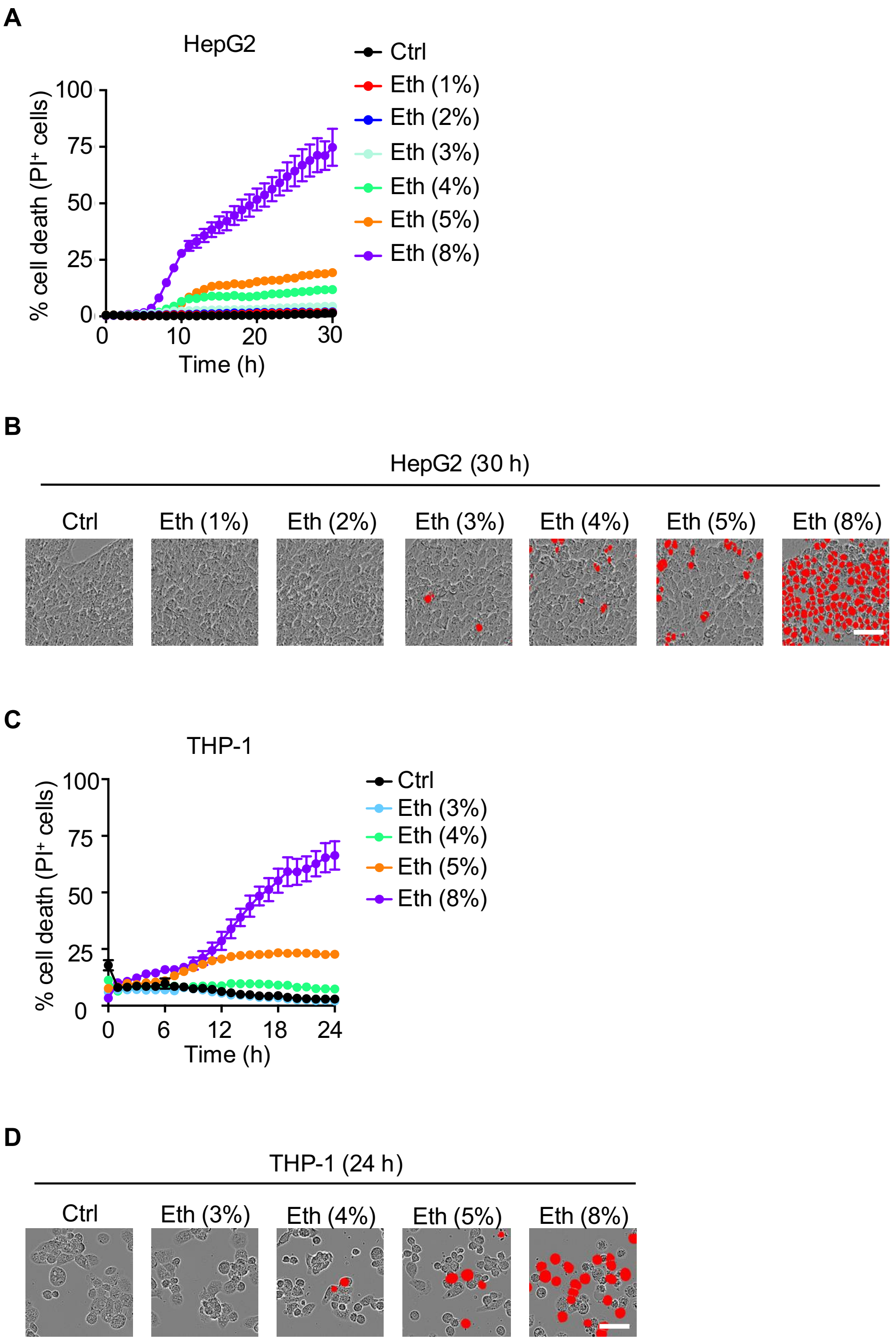


**Figure S2. Ethanol induces dose-dependent cell death in human liver and immune cells**
(**A**) Real-time cell death analysis of HepG2 cells left unstimulated in media (control; Ctrl) or treated with increasing concentrations of ethanol. (**B**) Representative images of HepG2 cells showing cell death 30 h post-ethanol treatment. (**C**) Real-time cell death analysis of human THP-1 cells left unstimulated in media (control; Ctrl) or treated with increasing concentrations of ethanol. **(D)** Representative images of THP-1 cells showing cell death 24 h post-ethanol treatment. The data are representative of at least three independent experiments. Scale bar, 50 μm in (**B and D**). The data are represented as the mean ± SEM in (**A and C**). Eth indicates ethanol, and Ctrl indicates control.

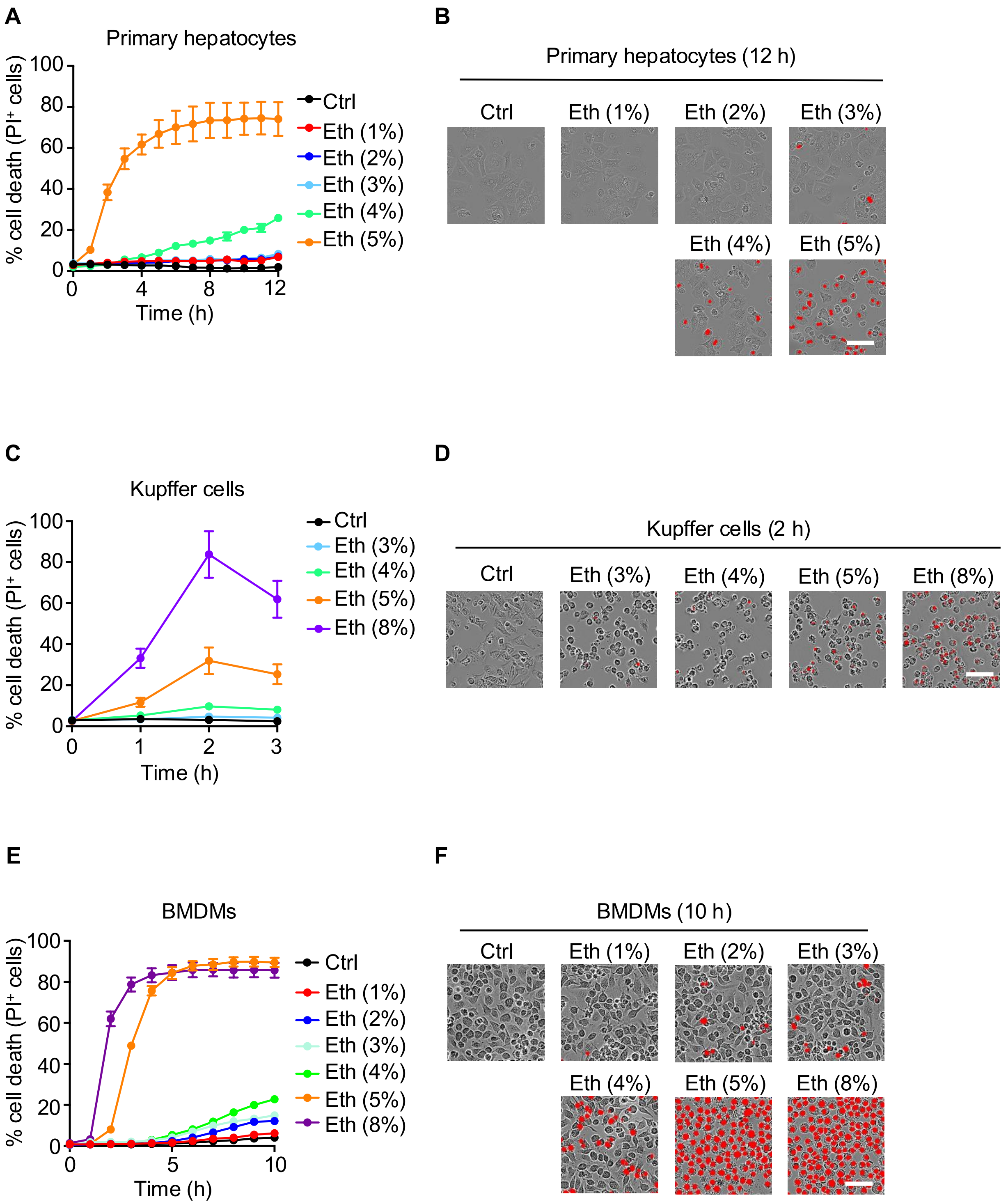


**Figure S3. Ethanol induces dose-dependent cell death in murine liver and immune cells**
(**A**) Real-time cell death analysis of mouse primary hepatocytes left unstimulated in media (control; Ctrl) or treated with increasing concentrations of ethanol. (**B**) Representative images of primary hepatocytes showing cell death 12 h post-ethanol treatment. (**C**) Real-time cell death analysis of Kupffer cells left unstimulated in media (control; Ctrl) or treated with increasing concentrations of ethanol. **(D)** Representative images of Kupffer cells showing cell death 2 h post-ethanol treatment. (**E**) Real-time cell death analysis of bone marrow-derived macrophages (BMDMs) left

unstimulated in media (control; Ctrl) or treated with increasing concentrations of ethanol. (**F**) Representative images of BMDMs showing cell death 10 h post-ethanol treatment. The data are representative of at least three independent experiments. Scale bar, 50 μm in (**B, D and F**). The data are represented as the mean ± SEM in (**A, C, and E**). Eth indicates ethanol, and Ctrl indicates control.

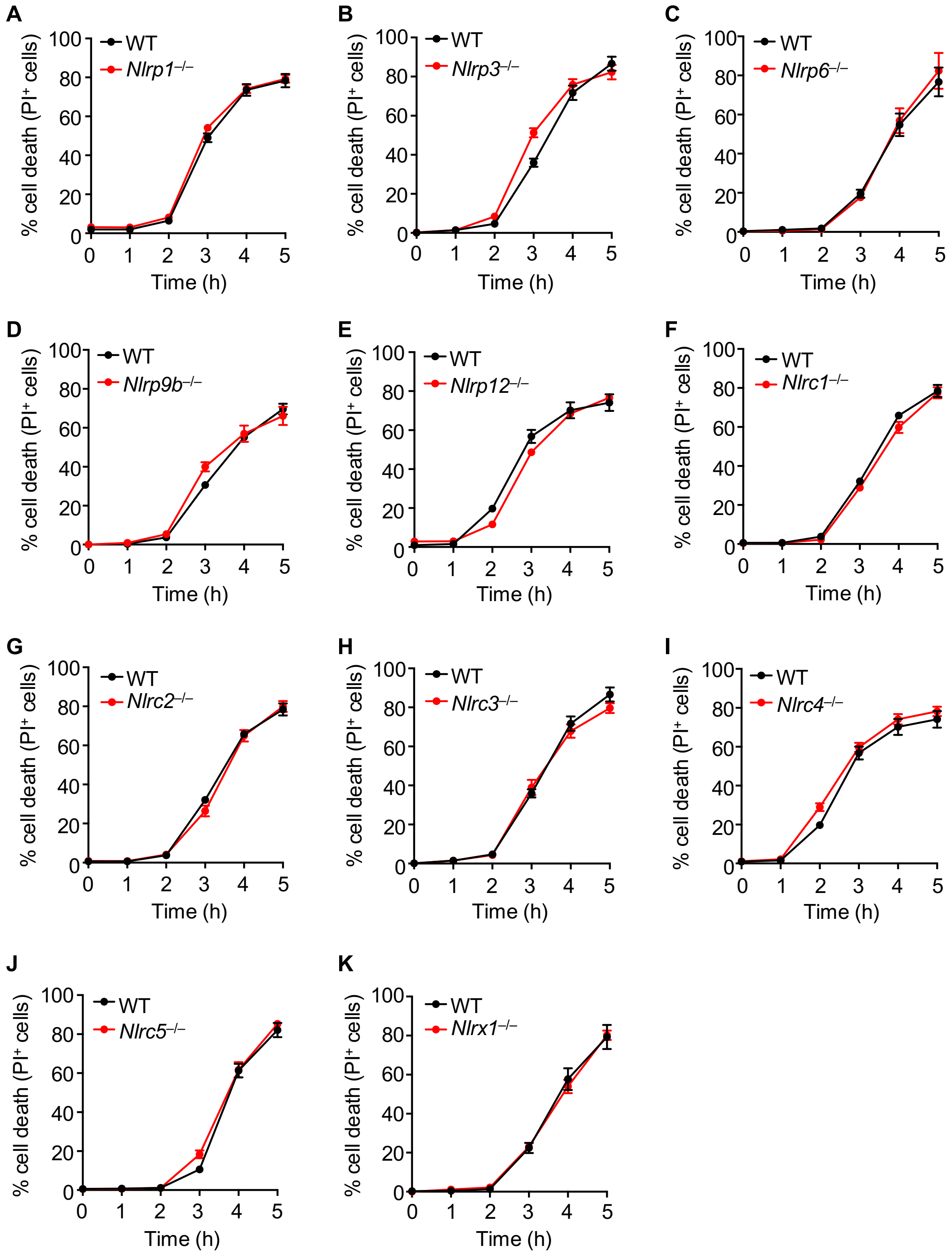


**Figure S4. NLRs do not contribute to ethanol-induced cell death**

(**A–K**) Real-time analysis of cell death in wild type (WT) versus *Nlrp1*$^{-/-}$ (**A**), *Nlrp3*$^{-/-}$ (**B**), *Nlrp6*$^{-/-}$ (**C**), *Nlrp9b*$^{-/-}$ (**D**), *Nlrp12*$^{-/-}$ (**E**), *Nlrc1*$^{-/-}$ (**F**), *Nlrc2*$^{-/-}$ (**G**), *Nlrc3*$^{-/-}$ (**H**), *Nlrc4*$^{-/-}$ (**I**), *Nlrc5*$^{-/-}$ (**J**), and *Nlrx1*$^{-/-}$ (**K**) bone marrow-derived macrophages (BMDMs) stimulated with ethanol. The data are representative of at least three independent experiments. The data are represented as mean ± SEM in (**A–K**). The analysis was performed using the two-way ANOVA (or mixed model) in (**A–K**).

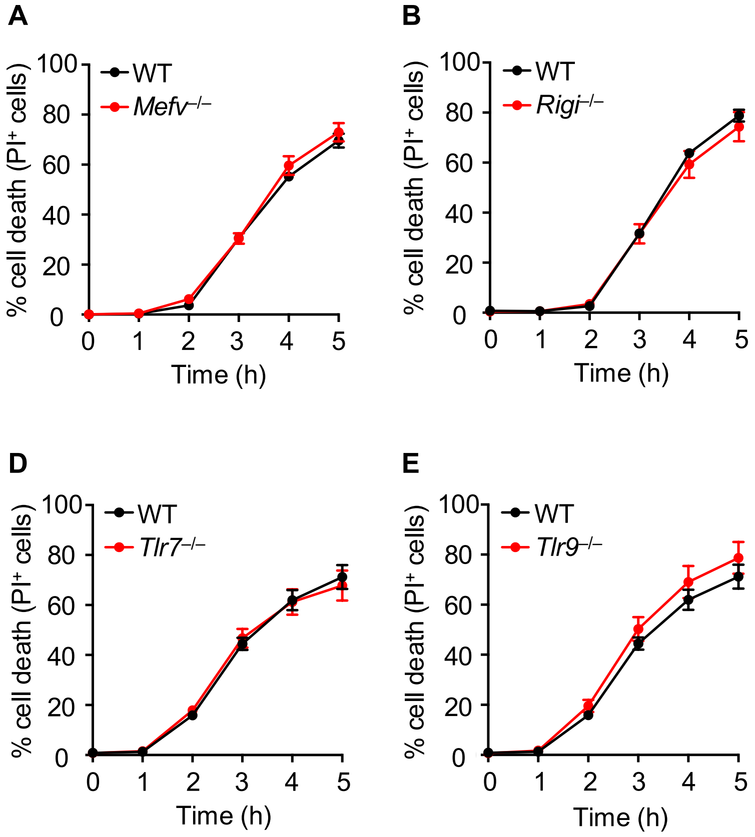


**Figure S5. Pyrin and RNA sensors do not contribute to ethanol-induced cell death**
(**A–F**) Real-time analysis of cell death in wild type (WT) versus *Mefv*$^{-/-}$ (**A**), *Rigi*$^{-/-}$ (**B**), *Tlr3*$^{-/-}$ (**C**), *Tlr7*$^{-/-}$ (**D**), *Tlr9*$^{-/-}$ (**E**), and *Tlr3/7/9*$^{-/-}$ (**F**) bone marrow-derived macrophages (BMDMs) stimulated with ethanol. The data are representative of at least three independent experiments. The data are represented as the mean ± SEM in (**A–F**). The analysis was performed using the two-way ANOVA (or mixed model) in (**A–F**).

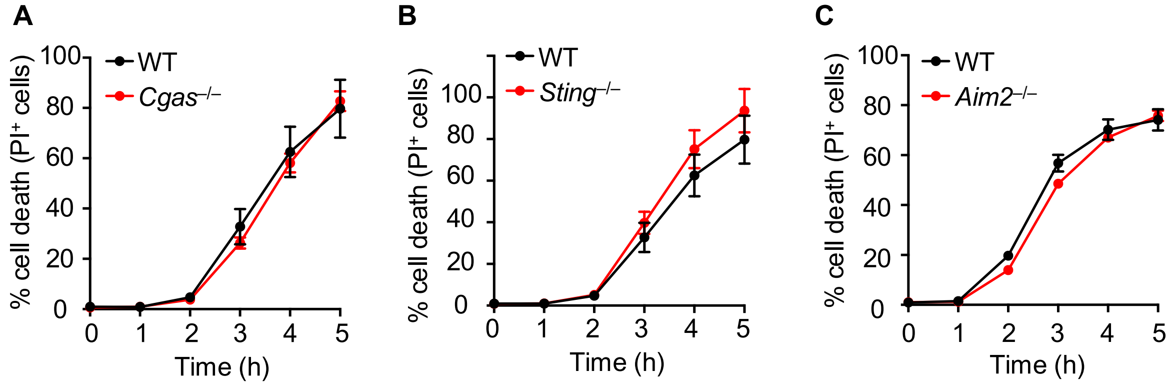


**Figure S6. cGAS/STING and AIM2 do not contribute to ethanol-induced cell death**
(**A–C**) Real-time analysis of cell death in wild type (WT) versus *Cgas*$^{-/-}$ (**A**), *Sting*$^{-/-}$ (**B**), and *Aim2*$^{-/-}$ (**C**) bone marrow-derived macrophages (BMDMs) stimulated with ethanol. The data are representative of at least three independent experiments. The data are represented as the mean ± SEM in (**A–C**). The analysis was performed using the two-way ANOVA (or mixed model) in (**A–C**).

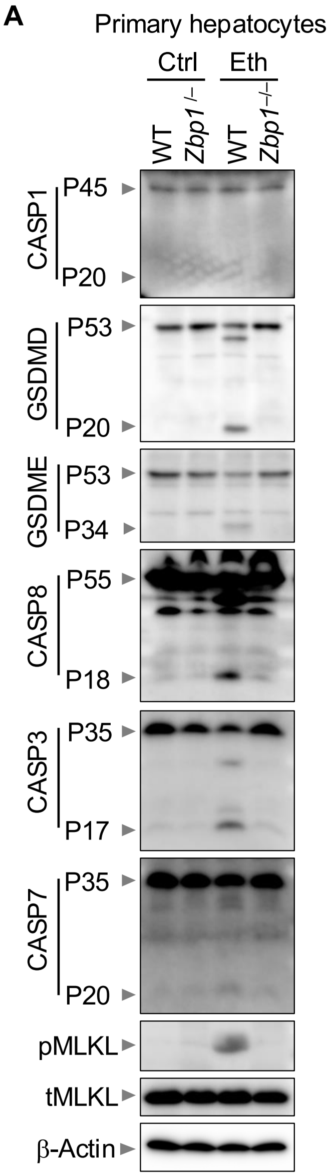


**Figure S7. ZBP1 drives ethanol-induced inflammatory cell death, PANoptosis, in primary hepatocytes**

(**A**) Immunoblot analysis of pro- (P45) and activated (P20) caspase-1 (CASP1); pro- (P53) and inactivated (P20) gasdermin D (GSDMD); pro- (P53) and activated (P34) gasdermin E (GSDME); pro- (P55) and cleaved (P18) caspase-8 (CASP8); pro- (P35) and cleaved (P17) caspase-3 (CASP3); pro- (P35) and cleaved (P20) caspase-7 (CASP7); phosphorylated mixed lineage kinase domain-like pseudokinase (pMLKL), and total MLKL (tMLKL) in wild type (WT) and *Zbp1*$^{-/-}$ primary hepatocytes left unstimulated in media (control; Ctrl) or stimulated with ethanol (Eth) for 12 h. β-Actin was used as the internal control. The data are representative of at least three independent experiments.

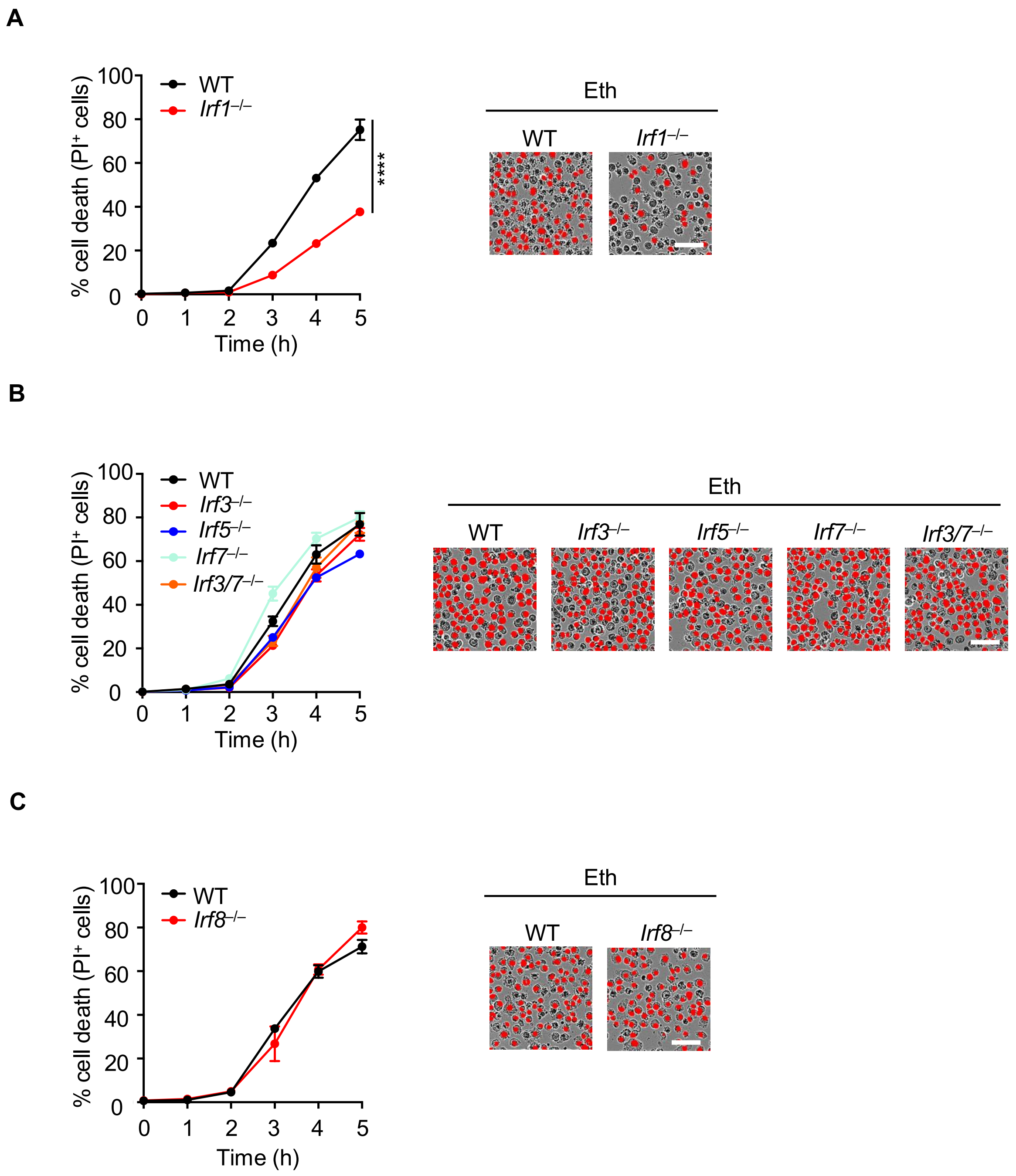


**Figure S8. IRF1, but not IRF3, IRF5, IRF7, or IRF8, contributes to ethanol-induced cell death** (**A–C**) Real-time cell death analysis in wild type (WT) versus *Irf1*$^{-/-}$ (**A**), *Irf3*$^{-/-}$, *Irf5*$^{-/-}$, *Irf7*$^{-/-}$, or *Irf3/7*$^{-/-}$ (**B**), or *Irf8*$^{-/-}$ (**C**) bone marrow-derived macrophages (BMDMs) stimulated with ethanol, and the representative images of cell death at 4 h post-ethanol treatment. The data are representative of at least three independent experiments. Scale bar, 50 µm. The data are represented as the mean ± SEM in (**A–C**). The analysis was performed using the two-way ANOVA (or mixed model) in (**A–C**). ****$P < 0.0001$.

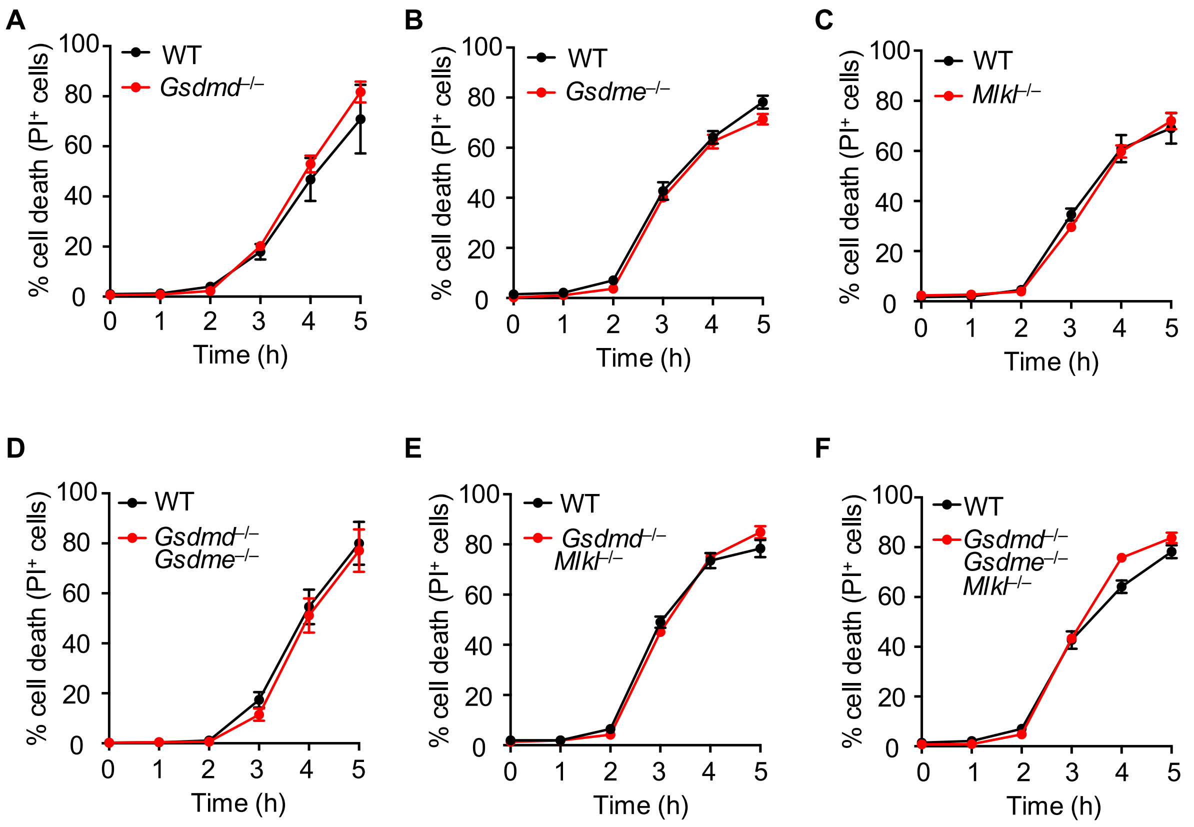


**Figure S9. Loss of GSDMD, GSDME, and MLKL does not block ethanol-induced cell death**
(**A–F**) Real-time analysis of cell death in wild type (WT) versus *Gsdmd*$^{-/-}$ (**A**), *Gsdme*$^{-/-}$ (**B**), *Mlkl*$^{-/-}$ (**C**), *Gsdmd*$^{-/-}$*Gsdme*$^{-/-}$ (**D**), *Gsdmd*$^{-/-}$*Mlkl*$^{-/-}$ (**E**), and *Gsdmd*$^{-/-}$*Gsdme*$^{-/-}$*Mlkl*$^{-/-}$ (**F**) bone marrow-derived macrophages (BMDMs) stimulated with ethanol. The data are representative of at least three independent experiments. The data are represented as the mean ± SEM in (**A–F**). The analysis was performed using the two-way ANOVA (or mixed model) in (**A–F**).

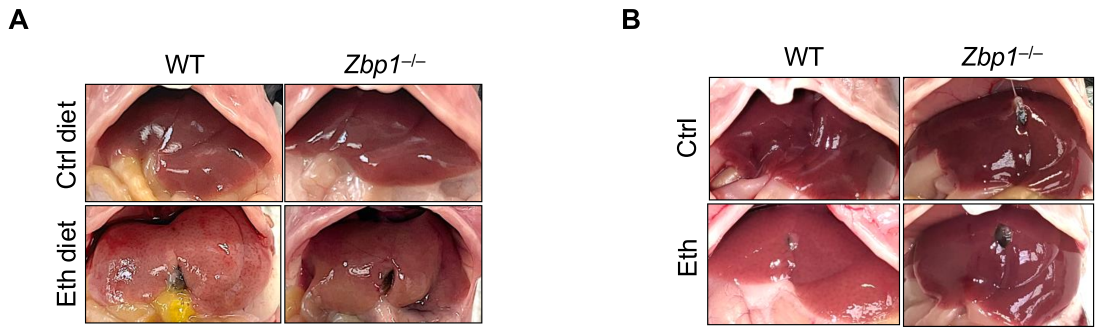


**Figure S10. ZBP1 contributes to the progression of ethanol-induced liver disease in mice**
(**A**) Representative images of livers from wild type (WT) and *Zbp1*$^{-/-}$ mice administered control and chronic-plus-binge ethanol diets. (**B**) Representative images of liver from WT and *Zbp1*$^{-/-}$ mice treated with or without ethanol in an acute alcoholic liver damage model. Eth indicates an ethanol diet or treatment, and Ctrl indicates a control diet or treatment.

**A**

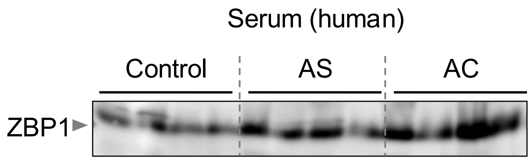


**Figure S11. Elevated ZBP1 in serum samples of patients with alcohol-associated liver disease**

(**A**) Immunoblot analysis of ZBP1 in 10 µl serum samples from individuals without alcohol-associated liver disease (control) and patients with alcoholic steatosis (AS) or alcoholic cirrhosis (AC).
